# NMDA Receptor Kinetics Drive Distinct Routes to Chaotic Firing in Pyramidal Neurons

**DOI:** 10.1101/2025.09.06.674628

**Authors:** Mehdi Borjkhani, Hadi Borjkhani, Morteza A. Sharif, Fariba Bahrami, Mahyar Janahmadi

## Abstract

Neuronal firing patterns emerge from complex interactions between intrinsic membrane properties and synaptic receptor dynamics. N-methyl-D-aspartate (NMDA) receptors critically shape calcium influx and synaptic plasticity through their voltage-dependent Mg^2+^ block and prolonged activation kinetics. We developed a Hodgkin-Huxley-type computational model incorporating NMDA, AMPA, and GABA receptor kinetics to investigate how NMDA receptor closing rates (*β*_*NMDA*_) and glutamatergic stimulation frequency control neuronal dynamics.

Systematic analysis of 2,942,093 inter-spike intervals across 1,961 parameter combinations revealed two mechanistically distinct pathways to firing irregularity. Pathway 1 involves rapid NMDA deactivation (*β*_*NMDA*_ *>* 0.06 ms^*−*1^) at elevated stimulation frequencies, producing deterministic chaos with compromised information encoding (entropy: 1.441 bits, mutual information: 0.185 bits). Pathway 2 results from slow NMDA deactivation (*β*_*NMDA*_ *<* 0.02 ms^*−*1^) under weak drive, creating irregularity through prolonged receptor activation and sustained calcium influx (entropy: 1.347 bits). An optimal kinetic window emerged at *β*_*NMDA*_ = 0.028 ms^*−*1^, maximizing information transfer (0.275 bits) while maintaining stable dynamics.

Entropy-Lyapunov correlation analysis confirmed deterministic chaos (r = 0.150, p ¡ 0.001). Frequency-dependent chaos onset thresholds demonstrated systematic erosion from 0.000 ms^*−*1^ at low frequencies to 0.150 ms^*−*1^ at high frequencies. GABAergic inhibition provided frequency-selective stabilization, expanding stable parameter space by 34.2

These findings establish NMDA receptor kinetics as fundamental controllers of cortical excitability and information processing. The dual-pathway framework provides mechanistic insights into addiction-related memory formation, where prolonged NMDA activation enables pathological plasticity, and visual processing disorders, where altered kinetics disrupt retinal function and cortical oscillatory balance. The identification of optimal kinetic windows and frequency-selective GABA modulation suggests therapeutic strategies targeting kinetically-specific interventions for neuropsychiatric disorders involving NMDA dysfunction.

## 1 Introduction

Information processing in the brain depends on precisely tuned interactions between intrinsic membrane conductances and synaptic receptor dynamics. NMDA receptors play a pivotal role in excitatory synaptic transmission, distinguished by their voltage-dependent Mg^2+^ block and relatively slow deactivation kinetics Kampa et al. (2004); Vargas-Caballero and Robinson (2004). These characteristics mediate extended calcium influx into postsynaptic neurons, influencing long-term potentiation (LTP), long-term depression (LTD), and other plasticity-related processes Lüscher and Malenka (2012). Beyond synaptic plasticity, NMDA receptors modulate neuronal firing stability and variability, with direct consequences for information encoding and transmission Harsch and Robinson (2000); Hunt and Castillo (2012).

A central question in computational neuroscience concerns how changes in NMDA receptor gating kinetics—particularly closing rates (*β*_*NMDA*_) and glutamate stimulation frequencies—transition neurons between stable periodic firing and irregular chaotic regimes Durstewitz and Gabriel (2007); Soudry and Meir (2012). While irregular spiking can degrade information transfer reliability, it may also increase dynamic range and encoding flexibility de Ruyter van Steveninck et al. (1997); Ermentrout et al. (2008). Despite extensive research on excitatory-inhibitory balance effects on neuronal dynamics, the detailed parameter space of NMDA receptor kinetics has received limited systematic investigation. The precise influence of NMDA receptor kinetics on plasticity induction, inter-spike interval (ISI) frequency band emergence, and their implications for single-cell information encoding remains incompletely understood.

The clinical significance of NMDA receptor kinetic alterations extends across multiple neuropsychiatric conditions. In schizophrenia, altered receptor kinetics may contribute to gamma oscillation abnormalities and cognitive deficits Coyle (2012); Gandal et al. (2012). Autism spectrum disorders involve NMDA-mediated excitation-inhibition imbalances affecting sensory processing and social cognition Rubenstein and Merzenich (2003); Lee et al. (2017). Alzheimer’s disease features progressive NMDA receptor dysfunction correlating with synaptic loss and memory impairment Wang and Reddy (2017); Snyder et al. (2005); Hynd et al. (2004). Chronic pain conditions exhibit altered NMDA kinetics in spinal cord circuits, contributing to central sensitization Latremoliere and Woolf (2009); Zhuo (2016).

In addiction neuroscience, chronic substance exposure alters NMDA receptor expression and kinetics, contributing to pathological synaptic plasticity underlying addiction-related memory formation and relapse vulnerability Kalivas (2009); Wolf (2016). Pathological memory formation represents a critical mechanism underlying addiction persistence and relapse vulnerability. Unlike normal learning and memory, drug-induced memories exhibit abnormal persistence, resistance to extinction, and heightened salience that can trigger craving and relapse even after prolonged abstinence Kalivas (2009); Wolf (2016). These pathological memories form through aberrant synaptic plasticity mechanisms involving dysregulated NMDA receptor function and downstream signaling cascades, particularly calcium-dependent processes such as CaMKII phosphorylation Lüscher and Malenka (2011); Pascoli et al. (2014).

Previous computational modeling has demonstrated that opioid exposure fundamentally alters synaptic plasticity mechanisms through disruption of calcium homeostasis and CaMKII phosphorylation dynamics Borjkhani et al. (2018a,b). These studies revealed that chronic drug exposure shifts the balance between LTP and LTD, favoring persistent synaptic modifications that encode drug-associated memories. NMDA receptors serve as critical gatekeepers in this process, with their kinetic properties determining calcium influx magnitude and persistence. Our research group has previously developed computational frameworks demonstrating how opioids induce pathological memory formation through theta rhythm generation during chronic consumption (Borjkhani et al., 2018c), and how cocaine disrupts action potential generation by reducing potassium currents (Borjkhani et al., 2022). These studies established the foundation for investigating drug-induced alterations in neural dynamics and synaptic plasticity mechanisms.

Evidence suggests NMDA receptor antagonists can disrupt drug memory reconsolidation and reduce relapse rates in preclinical studies Lee et al. (2006); Das et al. (2013). However, the precise relationship between NMDA receptor kinetic properties and pathological memory formation versus maintenance remains incompletely characterized. This knowledge gap represents a barrier to developing targeted therapeutic interventions that could selectively disrupt maladaptive memories while preserving normal memory function.

In visual neuroscience, NMDA receptors are essential for retinal ganglion cell function and cortical visual processing Manookin et al. (2010); Fox and Daw (1992). Receptor dysfunction contributes to glaucoma, diabetic retinopathy, and cortical visual impairment through altered firing patterns that disrupt normal visual processing Bai et al. (2013); Seki and Lipton (2008). In primary visual cortex, NMDA receptors mediate orientation selectivity refinement, ocular dominance plasticity, and contrast adaptation mechanisms Morishita and Hensch (2008); Hensch (2005). The kinetic properties of these receptors directly influence critical period timing and the capacity for experience-dependent plasticity throughout life Hensch (2005).

Recent evidence suggests altered NMDA kinetics contribute to visual processing deficits in neurodevelopmental disorders. In amblyopia, disrupted NMDA-dependent plasticity prevents normal binocular integration, while in autism spectrum disorders, altered excitation-inhibition balance affects visual motion processing and gamma oscillations Foss-Feig et al. (2013); Robertson and Baron-Cohen (2016). Age-related changes in NMDA receptor function may contribute to declining visual processing efficiency in older adults, with reduced NMDA-mediated plasticity affecting contrast sensitivity and temporal processing Hua et al. (2008).

Here, we develop and analyze a single-compartment Hodgkin-Huxley-type model of a pyramidal neuron incorporating sodium, potassium, and calcium currents, along with GABAergic, AMPA, and NMDA synaptic conductances. Building upon our previous computational investigations of opioid-induced memory formation (Borjkhani et al., 2018c) and cocaine effects on neuronal excitability (Borjkhani et al., 2022), we extend this framework to systematically examine NMDA receptor kinetic control of cortical dynamics. We include CaMKII phosphorylation dynamics as a biochemical pathway linking calcium transients to synaptic plasticity. By systematically varying *β*_*NMDA*_ and glutamatergic drive frequency, we evaluate how neuronal firing evolves through different oscillatory modes and assess impacts on spike-timing entropy, maximum Lyapunov exponent, and mutual information. We investigate how GABAergic inhibition modulates these transitions through fast inhibitory currents that control neuronal excitability and chaotic dynamics.

Our findings reveal dual pathways to firing irregularity and establish NMDA receptor kinetics as fundamental controllers of cortical excitability and information processing. These results provide mechanistic insights into addiction-related memory disorders and visual processing dysfunction, suggesting therapeutic strategies targeting NMDA receptor kinetic normalization and oscillatory pattern restoration.

## 2 Materials and Methods

Figure 1 provides an overview of our comprehensive computational approach to investigating NMDA receptor kinetics and their control over neuronal dynamics. The workflow encompasses four integrated stages: systematic model development and parameter space exploration, application of multiple dynamical analysis methods, identification of key mechanistic findings through the dual pathways framework, and translation to clinical applications. This systematic approach allows us to bridge molecular receptor properties with network-level dynamics and establish direct relevance to addiction neuroscience and visual processing disorders. The following sections detail each component of this workflow, beginning with the neuronal model architecture.

**Figure 1.**
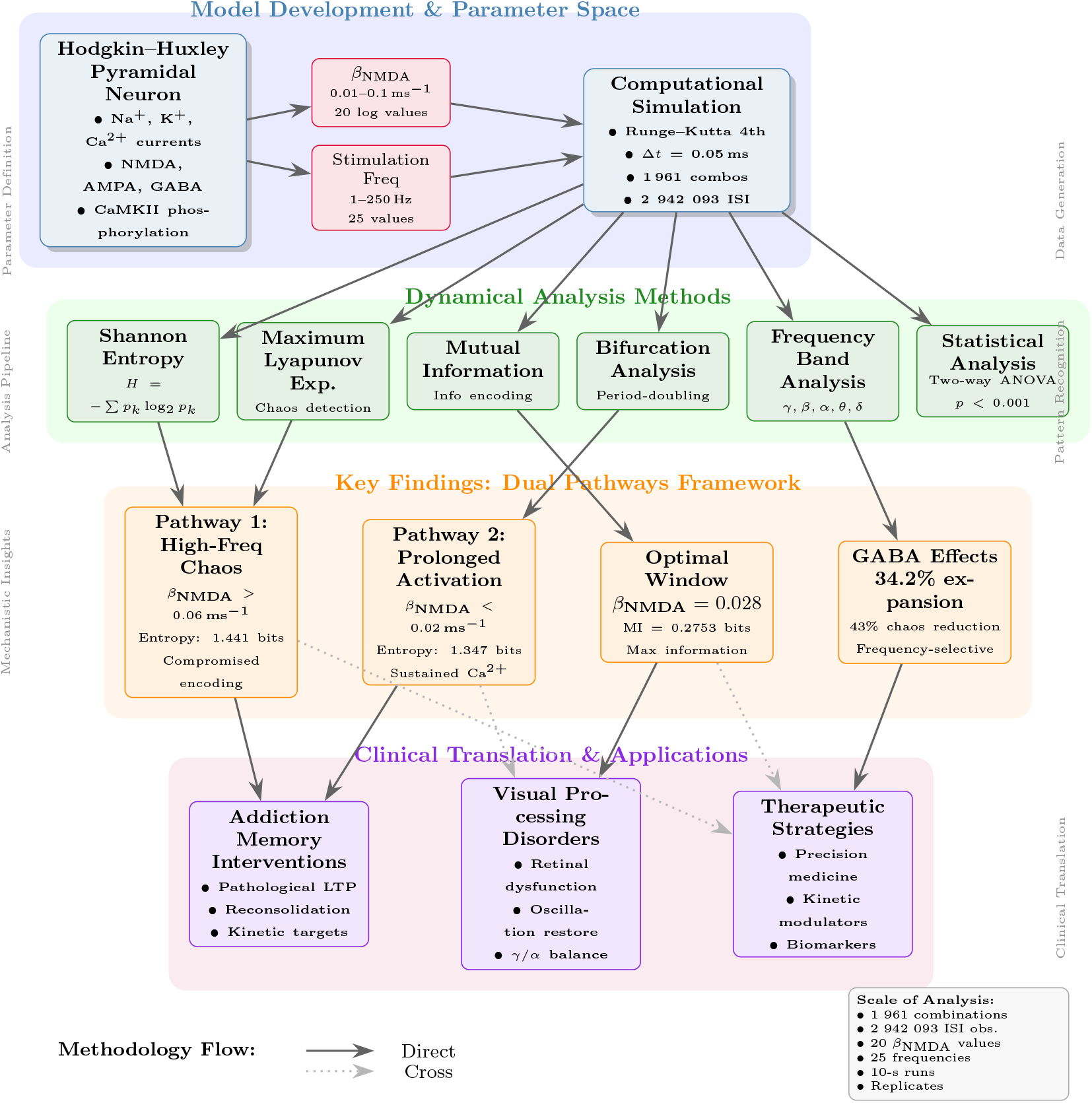
Comprehensive computational workflow for investigating NMDA receptor kinetics and neuronal dynamics. The study employed a systematic four-stage approach: (1) *Model Development & Parameter Space* Implementation of a detailed Hodgkin-Huxley pyramidal neuron model with systematic exploration of NMDA receptor closing rates (*β*_*NMDA*_: 0.01-0.1 ms^*−*1^, 20 values) and glutamatergic stimulation frequencies (1-250 Hz, 25 values). GABAergic modulation was tested at three levels (0, 25, 50 Hz), generating 1,500 parameter combinations (500 base × 3 GABA conditions). (2) *Dynamical Analysis Methods* Application of multiple complementary techniques including Shannon entropy analysis, maximum Lyapunov exponent calculation, mutual information quantification, bifurcation analysis, frequency band decomposition, and statistical validation. (3) *Key Findings: Dual Pathways Framework* Identification of two distinct routes to firing irregularity (Pathway 1: high-frequency chaos; Pathway 2: prolonged activation) and an optimal kinetic window for information encoding, alongside GABAergic modulation effects. (4) *Clinical Translation & Applications* -Direct relevance to addiction memory interventions, visual processing disorders, and therapeutic strategy development. The analysis encompassed 2,942,093 inter-spike interval observations across multiple computational replicates. Solid arrows indicate direct analyti-cal flow; dotted arrows represent cross-validation and mechanistic connections between findings and applications.

### 2.1 Neuronal Model Overview

For the simulation of a pyramidal neuron in the cortex, we employed a single-compartment HodgkinHuxley-type model based on Golomb et al. Golomb et al. (2006). The model was modified to incorporate glutamatergic and GABAergic receptors (NMDA, AMPA, and GABA receptors) alongside ionotropic channels, enabling responses to both excitatory and inhibitory neurotransmitters. Furthermore, the model incorporates CaMKII phosphorylation dynamics as a function of calcium concentration variations, providing a mechanistic link between synaptic activity and plasticity Borjkhani et al. (2018a). This computational framework allows investigation of NMDA receptor kinetics effects on neuronal excitability relevant to addiction-related plasticity mechanisms and visual processing disorders. Figure 2 shows the general structure of the modeled elements.

**Figure 2.**
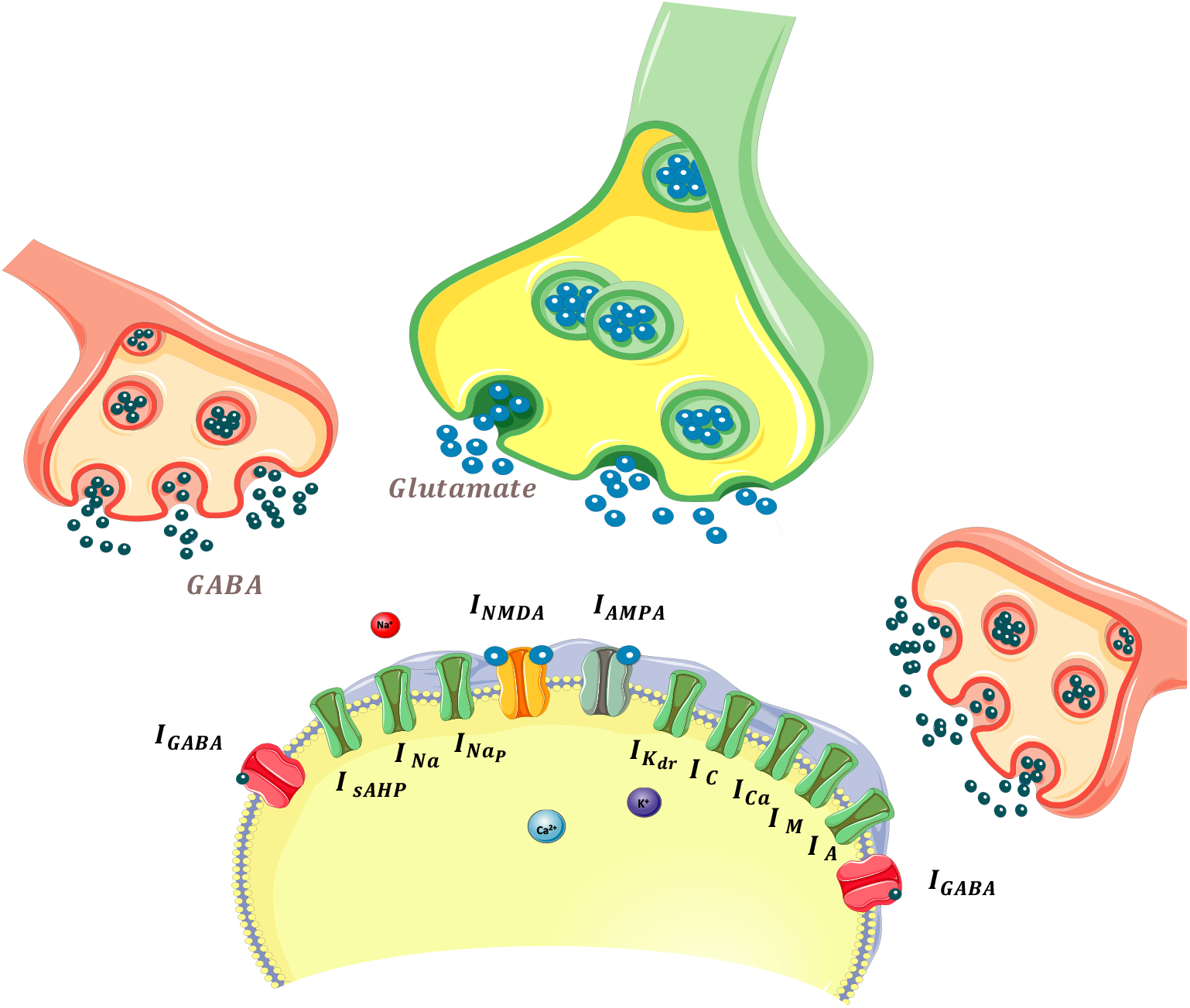
Synaptic inputs and ionic currents in the modeled neuron. This schematic illustrates the synaptic and intrinsic ionic currents included in the model. The total synaptic current consists of AMPA, NMDA, and GABAergic components, where AMPA and NMDA mediate excitatory transmission, with NMDA exhibiting a voltage-dependent Mg^2+^ block, while GABAergic currents provide inhibition. The intrinsic ionic currents include fast and persistent sodium (*I*_Na_, *I*_NaP_), various potassium currents regulating repolarization and adaptation (*I*_Kdr_, *I*_A_, *I*_M_), calciummediated currents (*I*_Ca_, *I*_C_), and slow afterhyperpolarization and leak currents (*I*_sAHP_, *I*_L_). These currents collectively shape neuronal excitability, synaptic integration, and firing dynamics.

#### 2.1.1 Model Validation and Verification

The model was validated against experimental data from pyramidal neurons in layers 2/3 of visual cortex Markram et al. (1997). Key validation metrics included: (1) resting potential (−65 ± 5 mV), (2) action potential amplitude (80-100 mV), (3) spike threshold (−45 ± 3 mV), and (4) adaptation ratio (0.6-0.8) during sustained current injection. Numerical integration was verified using analytical solutions for simplified cases and compared with NEURON simulator results (relative error *<* 0.1%).

### 2.2 Membrane Dynamics

The membrane potential (*V*) in the excitatory postsynaptic neuron is governed by the following differential equation:

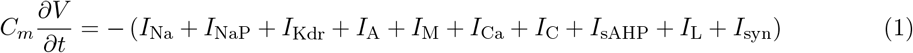

where the membrane capacitance *C*_*m*_ = 1 µF*/*cm^2^ and the total synaptic current is:

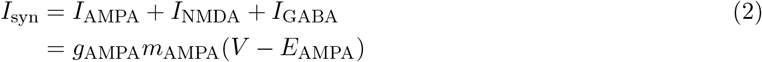

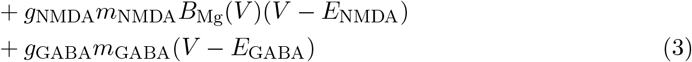

### 2.3 Intrinsic Ionic Currents

#### 2.3.1 Sodium Currents

The transient sodium current is described by:

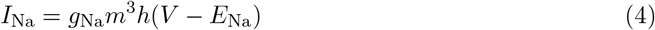

where *g*_Na_ = 35 mS/cm^2^ and *E*_Na_ = 55 mV is the sodium reversal potential. The activation and inactivation gating variables are denoted by *m* = *m*_∞_(*V*) and:

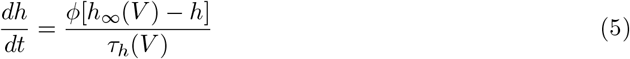

where *ϕ* = 5 is a temperature factor and the time constant is:

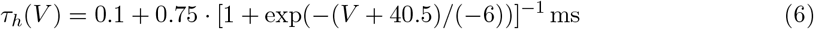

The persistent sodium current is:

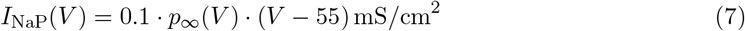

#### 2.3.2 Potassium Currents

The delayed rectifier potassium current is:

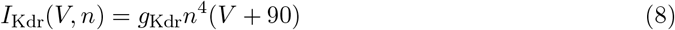

where *g*_Kdr_ = 6 mS/cm^2^ is the default conductance. The gating variable *n* follows:

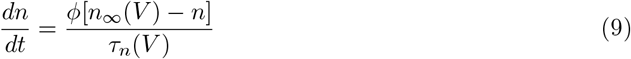

with time constant:

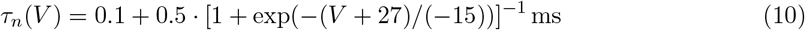

The A-type potassium current has the following dynamics:

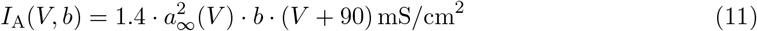

where *a* = *a*_∞_(*V*) and:

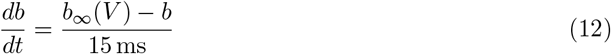

The muscarinic-sensitive potassium current is:

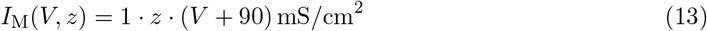

with gating variable dynamics:

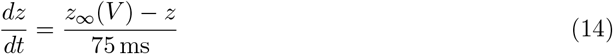

#### 2.3.3 Calcium Currents

The high-voltage calcium current is:

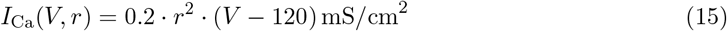

with gating variable:

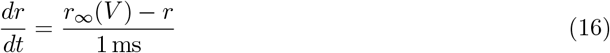

The fast calcium-activated potassium current is:

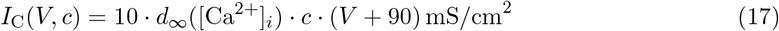

where [Ca^2+^]_*i*_ is the intracellular calcium concentration, and:

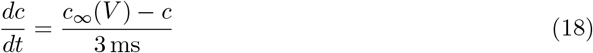

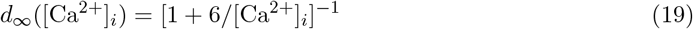

The slow calcium-activated potassium current (afterhyperpolarization) is:

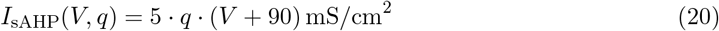

with gating variable dynamics:

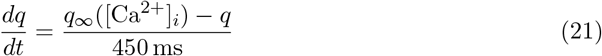

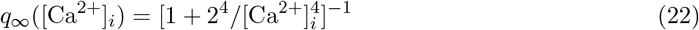

#### 2.3.4 Leak Current

The leak current is modeled as:

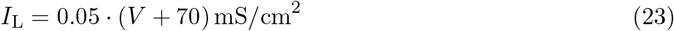

### 2.4 Calcium Dynamics

The intracellular calcium concentration evolves according to:

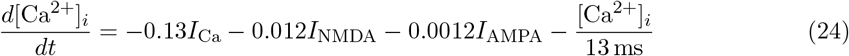

This equation accounts for calcium influx through voltage-gated calcium channels, NMDA receptors, and AMPA receptors (to a lesser extent), as well as calcium extrusion and buffering mechanisms with a time constant of 13 ms.

### 2.5 Synaptic Currents

#### 2.5.1 AMPA Receptors

The AMPA-mediated current is calculated as:

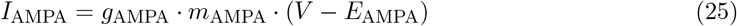

where *g*_AMPA_ = 0.5 nS and *E*_AMPA_ = 0 mV. The gating variable *m*_AMPA_ follows:

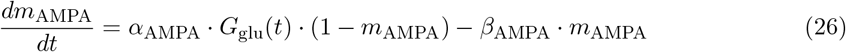

where *α*_AMPA_ = 1.1 mM^*−*1^ms^*−*1^ and *β*_AMPA_ = 0.67 ms^*−*1^ are the opening and closing rates, respectively. *G*_glu_(*t*) represents the time-varying glutamate concentration that activates the receptor.

#### 2.5.2 NMDA Receptors

The NMDA current is modeled as:

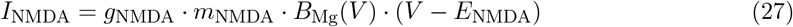

where *g*_NMDA_ = 0.5 nS and *E*_NMDA_ = 0 mV. The voltage-dependent Mg^2+^ block is:

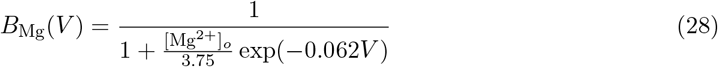

where [Mg^2+^]_*o*_ = 2 mM is the extracellular magnesium concentration.

The NMDA receptor gating variable follows:

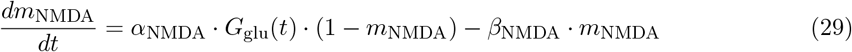

where *α*_NMDA_ = 0.14 mM^*−*1^ms^*−*1^ is the opening rate and *β*_NMDA_ is the systematically varied closing rate (range: 0.01–0.1 ms^*−*1^). This parameter represents the key experimental variable in our study, as alterations in NMDA receptor kinetics are implicated in both addiction-related plasticity and visual processing disorders.

#### 2.5.3 GABA Receptors

The GABAergic inhibitory current is:

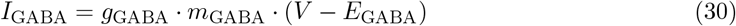

where *g*_GABA_ = 1.0 nS and *E*_GABA_ = −70 mV.

The GABA receptor gating variable follows:

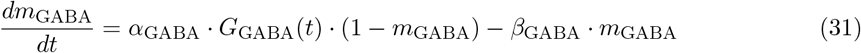

where *α*_GABA_ = 2.0 mM^*−*1^ms^*−*1^ and *β*_GABA_ = 1.5 ms^*−*1^.

### 2.6 Activation Functions

All steady-state activation and inactivation functions follow the standard Boltzmann form:

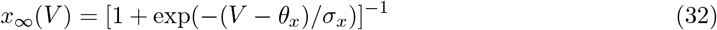

where *x* can be replaced by *m, h, n, a, b, z, p, r*, or *c*. The parameters are summarized in Table 1.

**Table 1.**
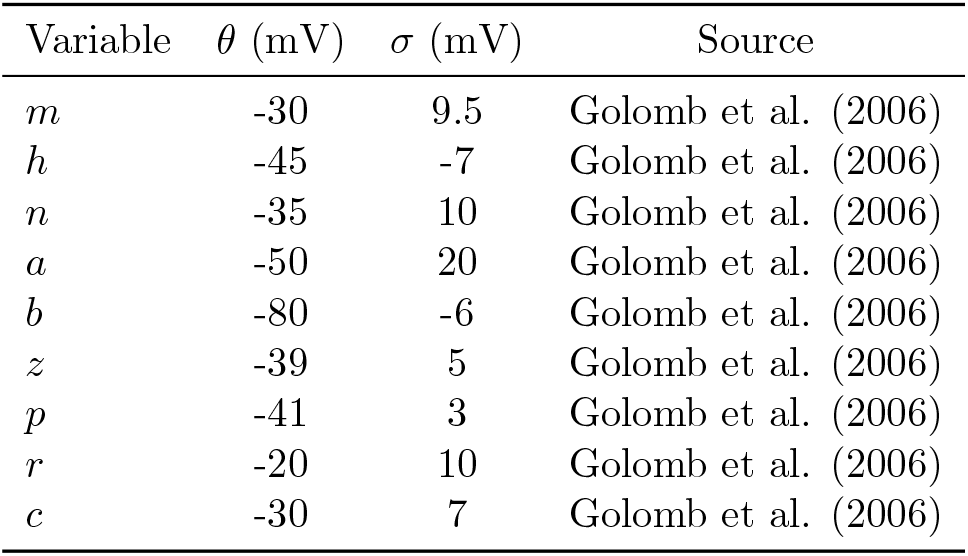

### 2.7 CaMKII Phosphorylation Dynamics

Postsynaptic Ca^2+^ concentration variations lead to CaMKII phosphorylation, which is governed by equations adapted from Borjkhani et al. Borjkhani et al. (2018a,b) and Zhabotinsky Zhabotinsky (2000):

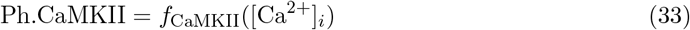

The detailed phosphorylation cascade involves 10 differential equations governing the concentrations of i-fold phosphorylated CaMKII (*P*_*i*_, where *i* = 0, 1, …, 10). The system includes:

### Phosphorylation rates

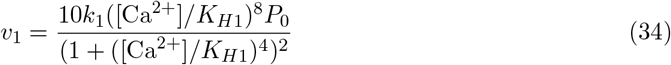

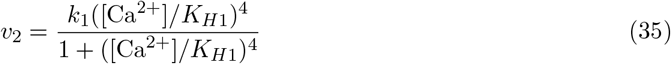

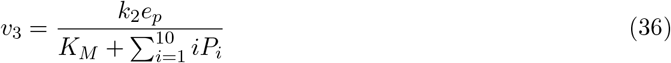

where *k*_1_ = 0.5 s^*−*1^ is the I1-dependent regulation rate of PP1, *K*_*H*1_ = 4 *µ*M is the Hill constant of CaMKII for calcium activation, *K*_*M*_ = 20 *µ*M and *k*_2_ = 10 s^*−*1^ are the Michaelis and catalytic constants, respectively.

### Additional parameters

- *e*_*p*_: PP1 concentration not bound to I1P (active protein phosphatase)
- *e*_*p*0_ = 0.1 *µ*M: total PP1 concentration
- *I*_0_ = 0.1 *µ*M: free I1 concentration
- *k*_3_ = 1 *µ*M^*−*1^s^*−*1^ and *k*_4_ = 10^*−*3^ s^*−*1^: association and dissociation rate constants of PP1-I1P complex
- *v*_CaN_ = 2 s^*−*1^: rate of I1P dephosphorylation due to calcineurin
- *v*_PKA_ = 0.45 *µ*M/s: phosphorylation rate of I1 due to PKA
- *K*_*H*2_ = 0.7 *µ*M: calcium activation Hill constant of calcineurin

The total phosphorylated CaMKII is:

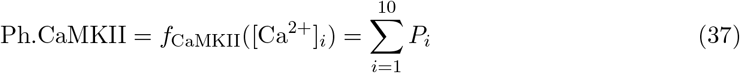

CaMKII phosphorylation levels were recorded at each time step and correlated with ISI patterns to investigate plasticity-related dynamics relevant to addiction and visual processing mechanisms.

### 2.8 Computational Implementation

#### 2.8.1 Software and Hardware Environment

All simulations were implemented in Python 3.8.10 using NumPy 1.21.0, SciPy 1.7.0, and Matplotlib 3.4.2. Computations were performed on a high-performance computing cluster with Intel Xeon Gold 6248 processors (2.5 GHz, 40 cores) and 192 GB RAM. Parallel processing utilized the multiprocessing library with 20 worker processes. Total computational time was approximately 480 CPU-hours.

#### 2.8.2 Numerical Integration and Simulation Parameters

All simulations employed fourth-order Runge-Kutta integration with Δ*t* = 0.05 ms, chosen based on convergence analysis showing *<* 0.1% error compared to Δ*t* = 0.01 ms. Simulation duration was 10 seconds with the first 2 seconds discarded to eliminate transients, determined from pilot studies showing equilibration within 1.5 seconds.

Glutamatergic stimulation consisted of 5 ms rectangular pulses with amplitude *G*_glu_ = 1 *µ*M and frequencies ranging from 1–250 Hz (25 logarithmically spaced values). GABAergic stimulation, when present, provided continuous background inhibition at three levels: 0 Hz (control), 25 Hz, and 50 Hz with *G*_GABA_ = 0.5 *µ*M.

#### 2.8.3 Parameter Space Exploration

We systematically varied *β*_NMDA_ across 20 logarithmically spaced values (0.01–0.1 ms^*−*1^) and stimulation frequency across 25 values (1–250 Hz), combined with 3 GABAergic conditions, generating 1,500 unique parameter combinations (20 × 25 × 3). Each combination was replicated *n* = 10 times with different random seeds (using numpy.random with seeds 0–9), totaling 15,000 simulations and yielding 2,942,093 ISI observations after quality control.

### 2.9 Data Analysis Pipeline

#### 2.9.1 Spike Detection and Quality Control

Action potentials were detected using a dual-threshold algorithm: initial detection at *V* = 0 mV, confirmation at *V* = 20 mV, with a 2 ms absolute refractory period. Inter-spike intervals were calculated as:

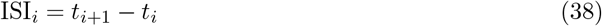

Quality control procedures excluded: (1) physiologically implausible ISIs (*<* 2 ms or *>* 2000 ms), (2) simulations with *<* 10 spikes, and (3) simulations showing numerical instability (voltage excursions *>* 200 mV). This resulted in exclusion of 1.2% of the total dataset.

#### 2.9.2 Dynamical Analysis Measures

##### Shannon Entropy

ISI variability was quantified using 20 equal-width bins based on Sturges’ rule for the dataset size:

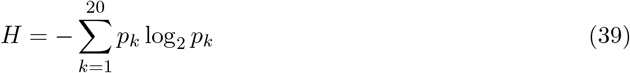

where *p*_*k*_ represents the probability of the *k*-th ISI bin. Bin width was determined individually for each parameter combination to ensure adequate sampling.

##### Maximum Lyapunov Exponents

Estimated using the Wolf et al. (1985) algorithm Wolf et al. (1985) with embedding dimension *d* = 5, time delay *τ* = 5 ms (determined from first minimum of mutual information), and nearest neighbor radius *r* = 0.1 mV. Convergence was verified by requiring *<* 5% change over the final 25% of the time series. Positive values (*λ >* 0) indicate chaotic dynamics.

##### Mutual Information

Calculated between parameters and ISI patterns using:

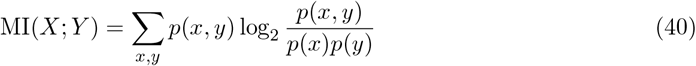

Bias correction was applied using the Miller-Madow estimator Miller (1955). Statistical significance was assessed using surrogate data generated by 1000 random permutations.

#### 2.9.3 Frequency Band Classification

ISIs were categorized into physiologically relevant bands based on inverse frequency relationships: gamma (7–33 ms, corresponding to 30–143 Hz), beta (33–77 ms, 13–30 Hz), alpha (77–125 ms, 8–13 Hz), theta (125–250 ms, 4–8 Hz), and delta (250–2000 ms, 0.5–4 Hz). Probability density functions were estimated using Gaussian kernel density estimation with Scott’s rule for bandwidth selection.

#### 2.9.4 Bifurcation and Phase Space Analysis

Bifurcation diagrams plotted steady-state ISI values against *β*_NMDA_ after 2000 ms equilibration. Local maxima and minima were identified using a peak-finding algorithm with minimum prominence of 5% of the dynamic range. Phase portraits were constructed by plotting membrane voltage *V* (*t*) versus its numerical derivative *dV/dt* calculated using central differences.

#### 2.9.5 Statistical Analysis and Validation

Results are presented as mean ± SEM across *n* = 10 replications. Statistical significance was assessed using two-way ANOVA (*α* = 0.05) with Bonferroni correction for multiple comparisons. Effect sizes are reported as partial eta-squared 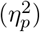. Assumptions were tested using Shapiro-Wilk tests for normality and Levene’s test for homoscedasticity.

Cross-validation was performed using 5-fold temporal splitting to assess stability of dynamical measures. Bootstrap confidence intervals (95%, *n* = 1000) were calculated for all summary statistics. Sensitivity analysis varied key parameters (integration timestep, bin numbers, embedding dimensions) by ±25% to ensure robustness of findings.

### 2.10 Model Limitations

The current model incorporates several simplifications: (1) single-compartment geometry neglecting dendritic processing, (2) simplified synaptic kinetics without vesicle depletion, (3) absence of network connectivity and population dynamics, and (4) deterministic framework excluding synaptic noise. These limitations were chosen to maintain computational tractability while preserving essential mechanisms relevant to NMDA receptor kinetics and neuronal excitability.

### 2.11 Reproducibility Statement

All simulation code and analysis scripts are available at https://github.com/borjkhani/Bifurcation_NMDA. Computational requirements include Python 3.8+ with specified dependencies and approx-imately 32 GB RAM for full parameter space exploration. Random seed handling ensures reproducible results, and detailed parameter files enable exact replication of all findings. Raw data and processed results are available upon reasonable request in accordance with institutional data sharing policies.

## 3 Results

### 3.1 Model Validation and Basic Neuronal Response

The computational model reproduced characteristic pyramidal neuron dynamics under controlled stimulation conditions (Figure 3). Glutamatergic inputs (1 *µ*M pulses) and continuous GABAergic background activity (0.5 *µ*M) generated distinct synaptic current profiles. NMDA currents exhibited slow kinetics with sustained activation, AMPA currents showed rapid transient responses, and GABA currents provided inhibitory modulation. Regular action potential firing occurred at 30.0 Hz with realistic membrane potential dynamics ranging from -75 mV to +50 mV. Intracellular calcium concentration ([Ca^2+^]_*i*_) oscillated between baseline and 0.858 *µ*M peaks (mean: 0.256 *µ*M), reflecting NMDA receptor-mediated calcium influx. CaMKII phosphorylation levels at *β*_NMDA_ = 0.0520 ms^*−*1^ demonstrated three distinct plasticity regimes: Long-Term Depression (LTD), metaplasticity transition zone, and Long-Term Potentiation (LTP) regions, with phosphorylation levels plotted on logarithmic scale over time.

**Figure 3.**
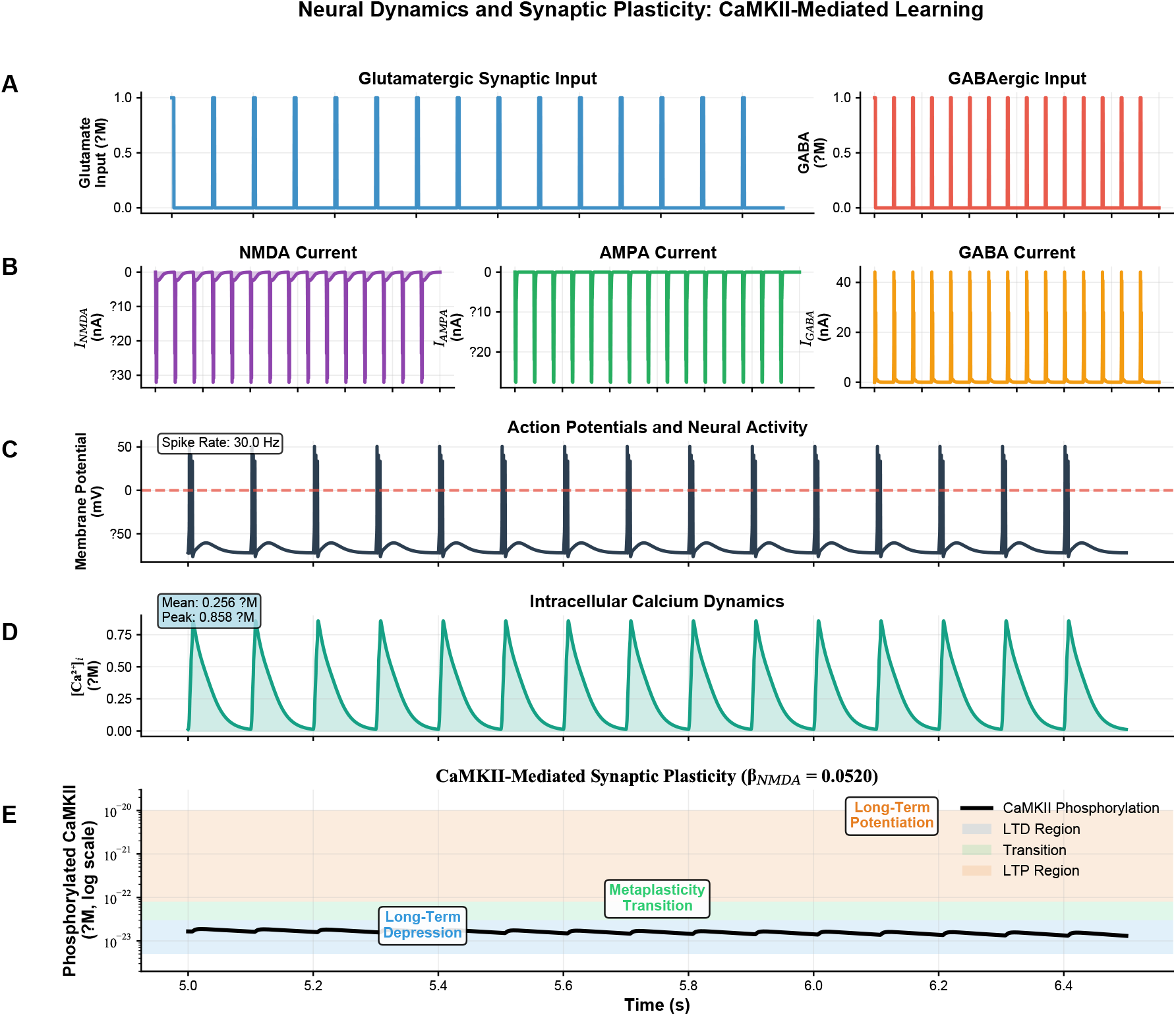
Neural dynamics and synaptic plasticity: CaMKII-mediated learning. (A) Synaptic input patterns: glutamatergic stimulation (left, 1 *µ*M pulses) and GABAergic inhibitory input (right, continuous background activity). (B) Synaptic currents showing NMDA-mediated (*I*_NMDA_), AMPA-mediated (*I*_AMPA_), and GABA-mediated (*I*_GABA_) responses during stimulation. (C) Action potential generation and neural activity with spike rate of 30.0 Hz, demonstrating realistic membrane potential dynamics. (D) Intracellular calcium dynamics ([Ca^2+^]_*i*_) with mean concentration of 0.256 *µ*M and peak values reaching 0.858 *µ*M, reflecting NMDA receptor-mediated calcium influx. (E) CaMKII phosphorylation dynamics (*β*_NMDA_ = 0.0520) showing distinct plasticity regimes: Long-Term Depression (LTD) region, metaplasticity transition zone, and Long-Term Potentiation (LTP) region, with CaMKII phosphorylation levels plotted on logarithmic scale over time.

### 3.2 Information-Theoretic Analysis Reveals Optimal NMDA Receptor Modulation

Information-theoretic analysis identified optimal NMDA receptor kinetics for neural coding (Figure 4). Mutual information between neural responses and stimuli peaked at *β*_NMDA_ = 0.028 ms^*−*1^ with maximum information transfer of 0.275 bits. Mutual information exhibited frequency-dependent characteristics, with highest values in the low-frequency range (0-30 Hz) and rapid decay at higher stimulation frequencies above 50 Hz. Neural response entropy increased with NMDA modulation strength, reaching a plateau at *β*_NMDA_ = 0.383 ms^*−*1^ with peak entropy of 1.81 bits. Entropy demonstrated frequency-dependent modulation with a prominent peak at 185 Hz (2.89 bits), followed by sharp decline at higher frequencies. Error bars represent standard deviation across multiple trials.

**Figure 4.**
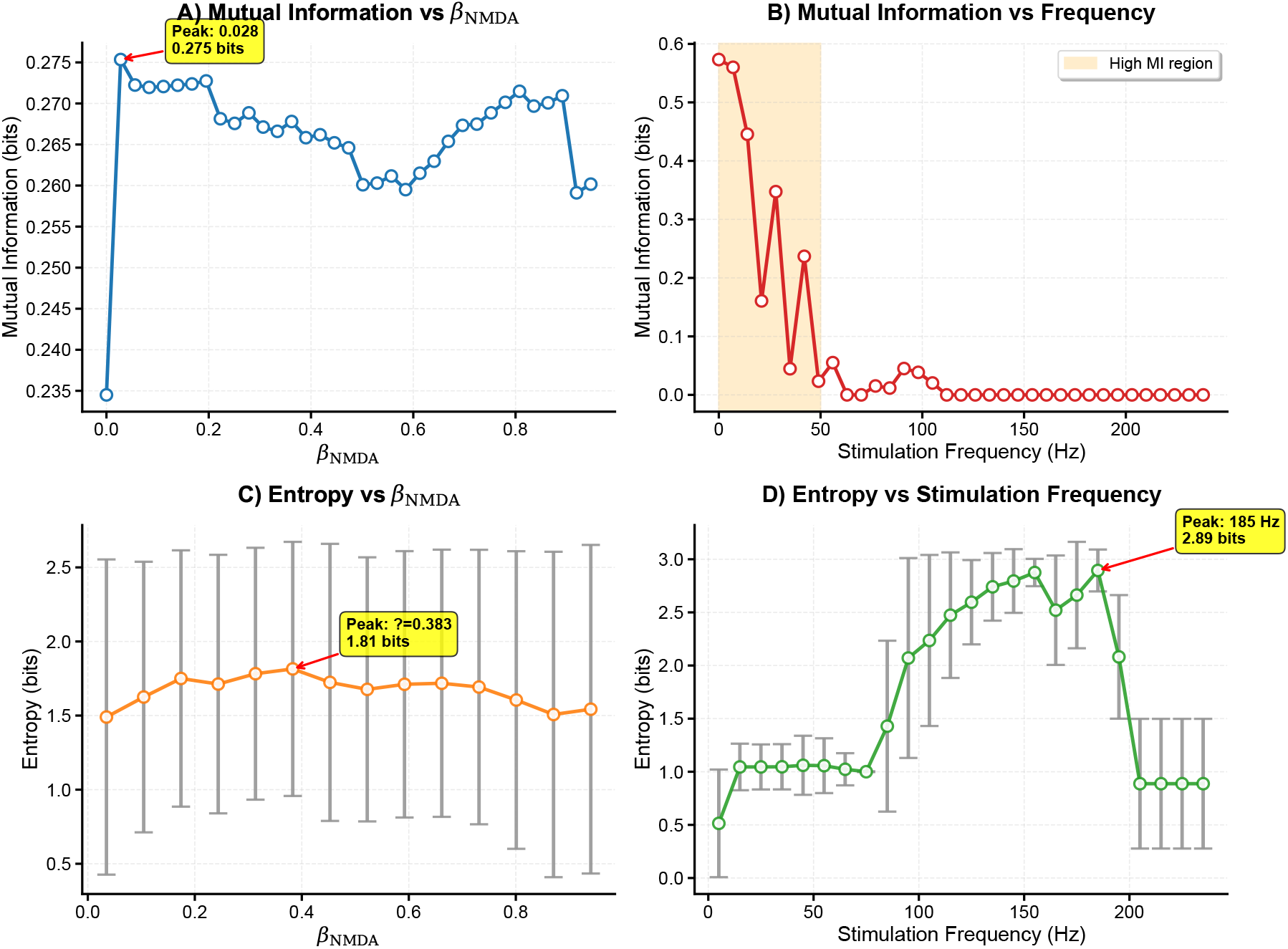
Information-theoretic analysis reveals optimal NMDA receptor modulation and frequency-dependent neural coding. (A) Mutual information between neural responses and stimuli as a function of NMDA receptor modulation strength (*β*_NMDA_). Peak information transfer occurs at *β*_NMDA_ = 0.028, corresponding to 0.275 bits. (B) Mutual information varies with stimulation frequency, showing maximal information content in the low-frequency range (0-30 Hz, shaded region) with rapid decay at higher frequencies. (C) Neural response entropy increases with NMDA modulation strength, reaching a plateau around *β*_NMDA_ = 0.4 (peak: 1.81 bits at *β*_NMDA_ = 0.383). Error bars represent standard deviation across trials. (D) Entropy exhibits frequencydependent modulation with a prominent peak at 185 Hz (2.89 bits), followed by a sharp decline. Error bars represent standard deviation. The analysis demonstrates that moderate NMDA receptor modulation optimizes information transfer while maintaining response diversity, with distinct coding regimes for low-frequency information transmission and high-frequency response entropy.

### 3.3 Bifurcation Analysis Reveals Period-Doubling Routes to Chaos

Detailed bifurcation analysis in the low-frequency regime revealed systematic transitions in firing dynamics (Figure 5). Entropy and maximum Lyapunov exponent heatmaps identified chaotic regions concentrated in low-frequency, low-*β*_NMDA_ parameter space. Phase portraits demonstrated progressive evolution from regular periodic firing (*β*_NMDA_ = 0.005 ms^*−*1^) through period-doubling cascades to chaotic dynamics (*β*_NMDA_ = 0.08 ms^*−*1^). Bifurcation diagrams showed period-doubling routes to chaos with pink regions indicating chaotic parameter ranges. ISI probability density functions revealed frequency band evolution from gamma (30-100 Hz) through beta (13-30 Hz) to alpha (8-13 Hz) distributions. CaMKII phosphorylation analysis across different *β*_NMDA_ values (0.007, 0.042, 0.082 ms^*−*1^) demonstrated distinct plasticity regimes: LTP region (high phosphorylation), transition zone, and LTD region (low phosphorylation).

**Figure 5.**
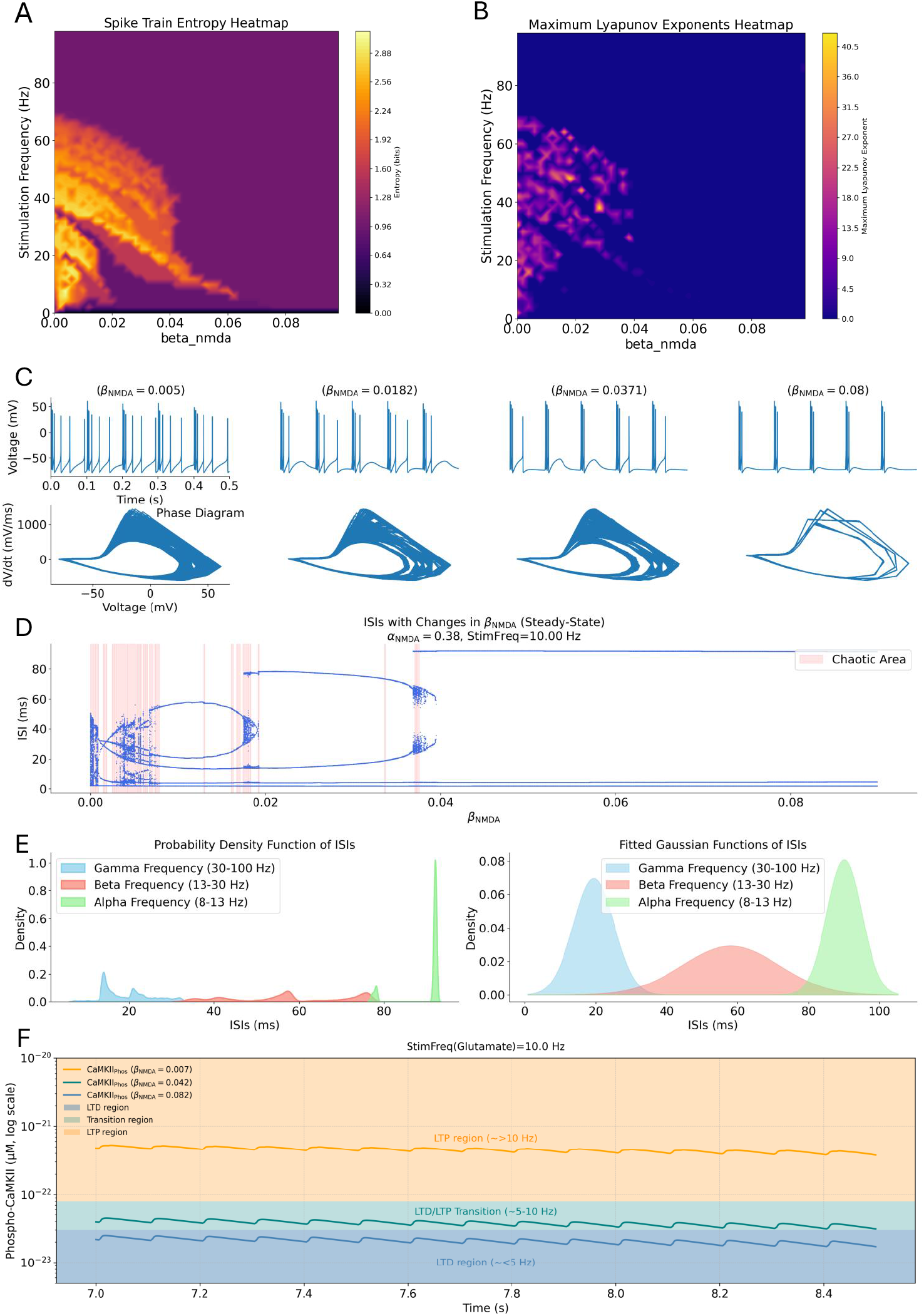
Bifurcation analysis and oscillatory dynamics. (A-B) Entropy and MLE heatmaps for low-frequency, low-*β*_NMDA_ parameter space. (C) Phase portraits showing transition from regular to chaotic firing. (D) Bifurcation diagram revealing period-doubling route to chaos (red regions indicate chaotic dynamics). (E) ISI probability density functions categorized by frequency bands. (F) CaMKII phosphorylation levels across plasticity regimes: LTP (orange), transition (green), LTD (blue).

### 3.4 Frequency-Dependent Neural Dynamics and Phase Space Evolution

Voltage traces and phase diagrams demonstrated systematic changes in neural dynamics across stimulation frequencies (Figure 6). At 2 Hz stimulation, neurons exhibited complex firing patterns with multiple *β*_NMDA_ values producing distinct voltage trajectories and elaborate phase space structures. At 5 Hz, moderate frequency stimulation showed earlier onset of irregular dynamics with simplified phase portraits. Higher stimulation frequencies produced progressive stabilization: 15 Hz demonstrated rapid convergence to regular firing patterns, while 50 Hz stimulation resulted in highly regular dynamics with compressed phase space trajectories. Phase diagrams revealed systematic evolution from complex multi-loop attractors at low frequencies to simple limit cycles at high frequencies. Voltage amplitudes ranged from -75 mV to +50 mV across all conditions, with firing patterns becoming increasingly predictable as stimulation frequency increased.

**Figure 6.**
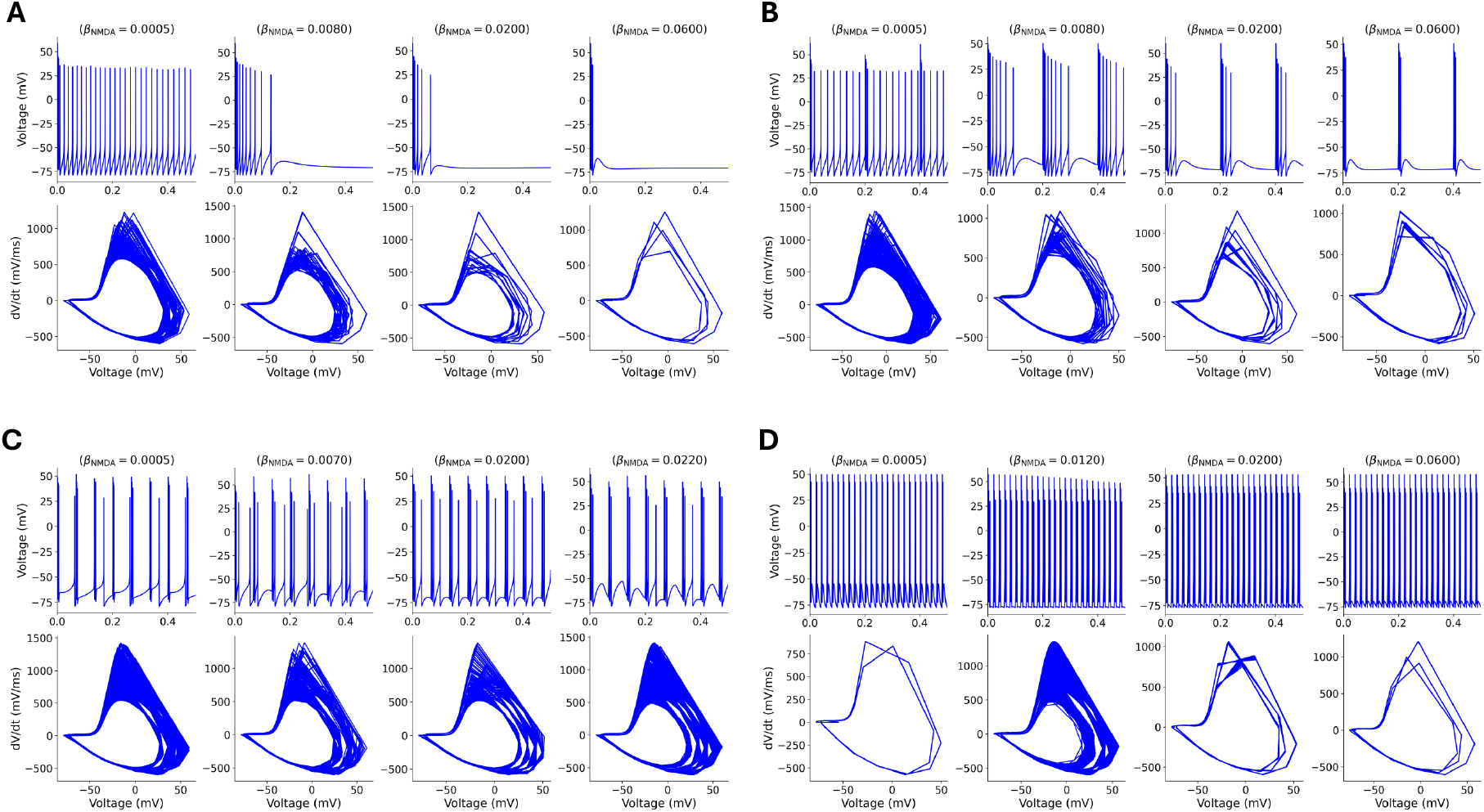
Neural dynamics across stimulation frequencies. Voltage traces and phase diagrams showing progressive destabilization from (A) 2 Hz stable dynamics with complex bifurcations, (B) 5 Hz moderate frequency showing earlier chaos onset, (C) 15 Hz rapid destabilization, to (D) 50 Hz immediate chaotic dynamics with highly irregular patterns.

### 3.5 Frequency-Dependent Chaos Threshold Erosion

Bifurcation diagrams revealed systematic erosion of chaos thresholds with increasing stimulation frequency (Figure 7). At 2 Hz stimulation, chaotic regions (pink shading) occupied 18 parameter ranges (2.0% of *β*_NMDA_ space), with complex bifurcation structures and ISI values ranging from 0-500 ms. At 5 Hz, chaotic regions decreased to 22 ranges (2.4% coverage) with ISI compression to 0-200 ms. Further frequency increases produced progressive stabilization: 10 Hz exhibited 25 chaotic regions (2.8% coverage) with ISI range 0-80 ms, while 15 Hz showed 20 regions (2.2% coverage) and ISI range 0-60 ms. At 50 Hz stimulation, chaos was nearly eliminated with only 5 regions (0.6% coverage) and tight ISI clustering around 15-18 ms. Bifurcation patterns evolved from complex multi-branch structures at low frequencies to simple periodic solutions at high frequencies.

**Figure 7.**
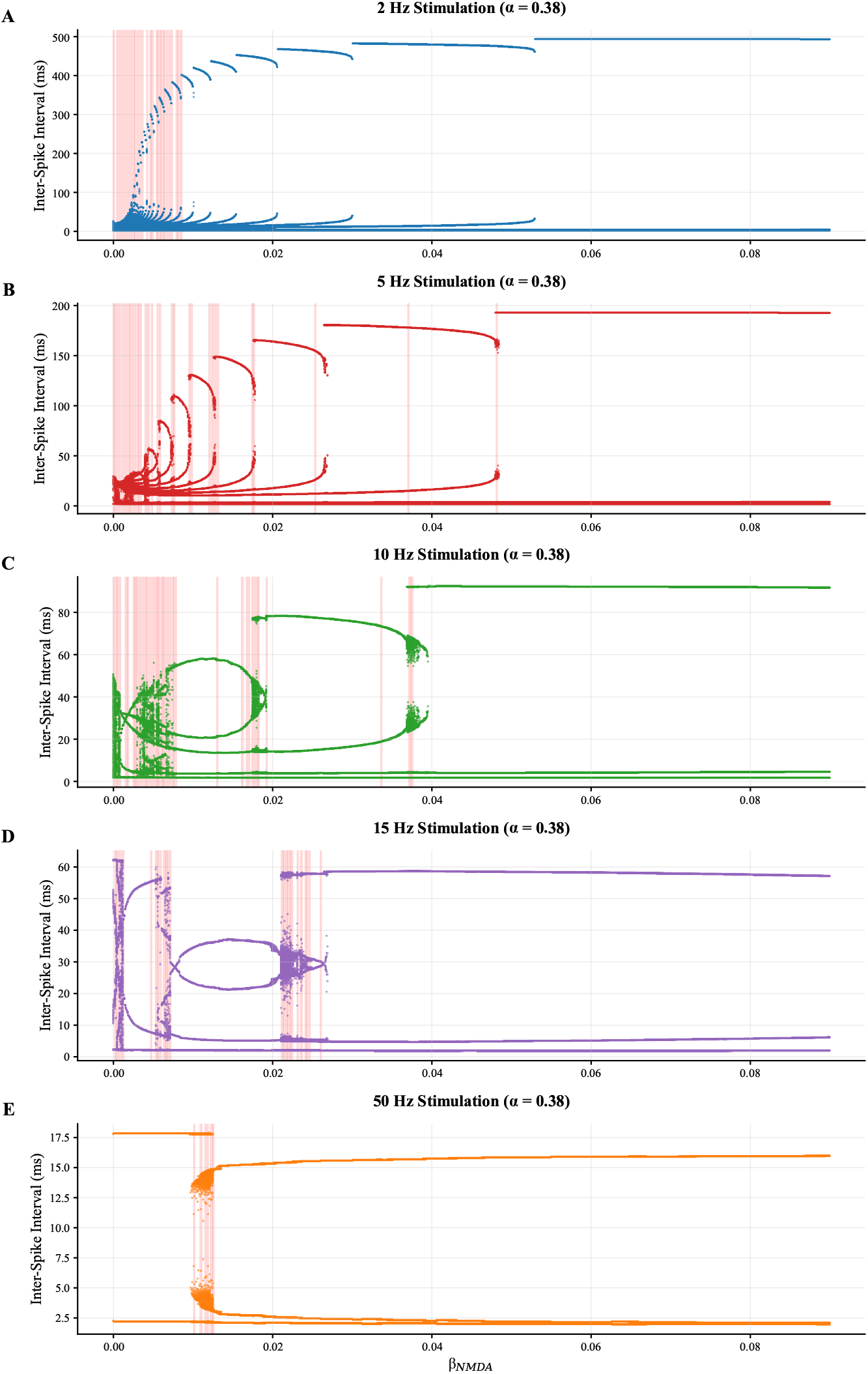
Frequency-dependent bifurcation landscapes and chaos onset thresholds. (A-E) Bifurcation diagrams showing inter-spike interval distributions versus *β*_NMDA_ across stimulation frequencies from 2 Hz to 50 Hz. Pink shading indicates chaotic regions. Progressive frequency increases demonstrate systematic erosion of stability thresholds: (A) 2 Hz exhibits complex bifurcations with chaos onset at *β*_NMDA_ ≈ 0.012 ms^*−*1^, (B) 5 Hz shows earlier destabilization, (C) 10 Hz displays limited chaotic windows, (D) 15 Hz31demonstrates compressed parameter ranges, and (E) 50 Hz maintains remarkable stability with tight gamma-frequency ISI clustering (15-18 ms) and minimal chaotic behavior.

### 3.6 Oscillatory Band Evolution and Spectral Consolidation

Frequency band analysis revealed systematic spectral evolution across stimulation rates (Figure 8). At 2 Hz stimulation, ISI distributions showed broad multi-band characteristics spanning gamma (30-100 Hz, blue), beta (13-30 Hz, red), alpha (8-13 Hz, green), theta (4-8 Hz, purple), and delta (0.5-4 Hz, yellow) frequency ranges. Empirical distributions demonstrated peak densities of 0.08 for beta band and 0.05 for alpha band. At 5 Hz stimulation, spectral content consolidated with reduced delta component and enhanced gamma representation. Progressive frequency increases produced systematic band elimination: 10 Hz stimulation showed dominant alpha band concentration (density = 1.0) with minimal beta and theta contributions. At 15 Hz, beta band dominance emerged (density = 0.5) with compressed alpha representation. At 50 Hz stimulation, complete gamma band concentration occurred (density = 3.5) with elimination of all slower frequency components. Gaussian mixture model fits confirmed progressive spectral narrowing from multi-component distributions to single-component gamma-range concentration.

**Figure 8.**
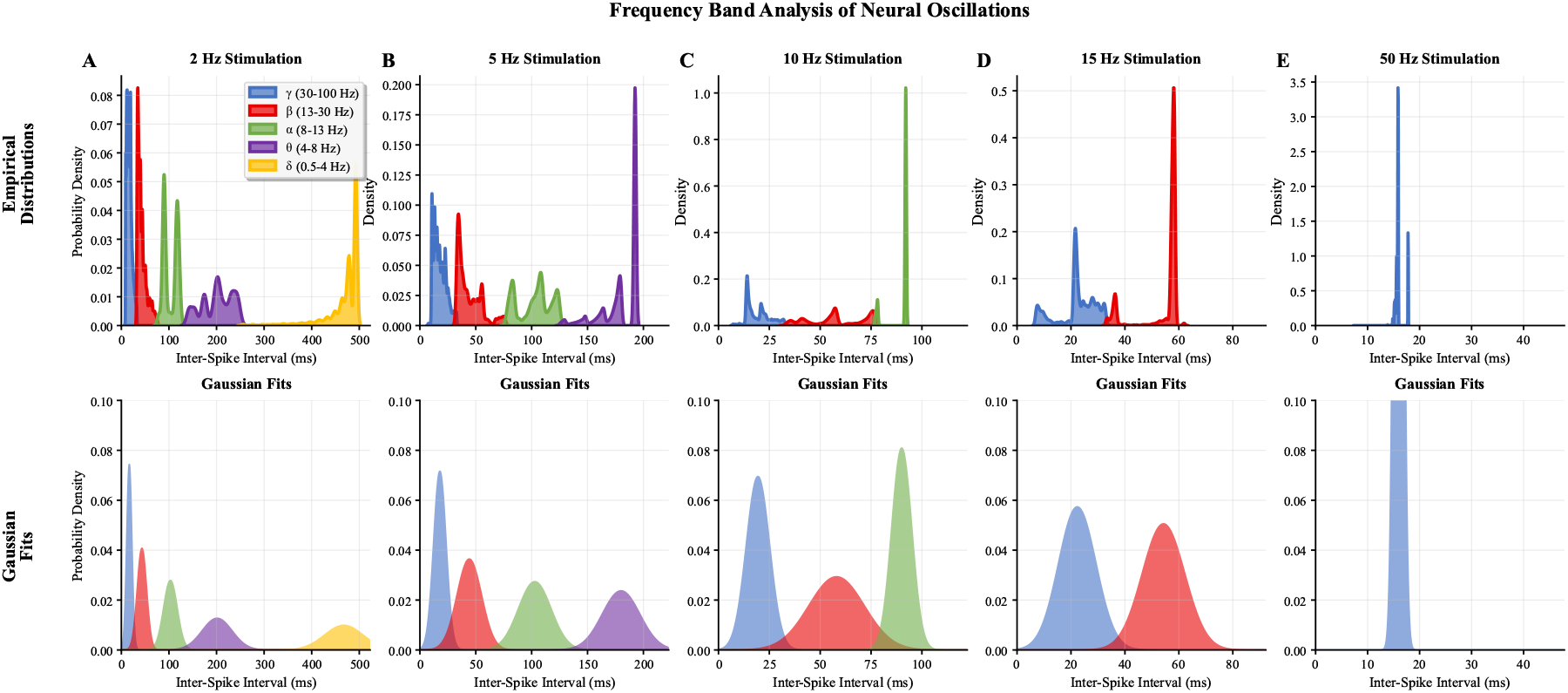
Frequency band analysis of neural oscillations across stimulation rates. (A-E) Top panels show empirical probability density distributions of inter-spike intervals decomposed into physiological frequency bands: gamma (30-100 Hz, blue), beta (13-30 Hz, red), alpha (8-13 Hz, green), theta (4-8 Hz, purple), and delta (0.5-4 Hz, yellow) across stimulation frequencies from 2 Hz to 50 Hz. Bottom panels display corresponding Gaussian mixture model fits. Progressive stimulation rate increases produce systematic spectral consolidation: (A) 2 Hz shows broad multi-band distributions, (B-D) intermediate frequencies (5-15 Hz) exhibit gradual gamma-band dominance, and (E) 50 Hz stimulation results in tight gamma-range concentration with minimal spectral diversity.

### 3.7 GABAergic Inhibition Provides Frequency-Selective Stabilization

Background GABAergic inhibition systematically modulated neural bifurcation dynamics across inhibition frequencies from 0-50 Hz (Figure 9). Control conditions (no GABA) exhibited extensive chaotic regions (pink shading) with complex bifurcation structures spanning ISI ranges from 0-100 ms. GABA inhibition at 2 Hz produced minimal stabilization effects, maintaining similar chaotic parameter coverage. Progressive GABA frequency increases demonstrated systematic stabilization: 5 Hz inhibition reduced chaotic regions and simplified bifurcation patterns, while 10 Hz further compressed chaotic parameter space. At 15 Hz and 20 Hz GABA inhibition, chaotic regions were substantially reduced with ISI ranges constrained to 20-80 ms. Complete chaos elimination occurred at 50 Hz GABA inhibition, producing stable dynamics with ISI clustering around 15-20 ms across the entire *β*_NMDA_ parameter range. Bifurcation structures evolved from complex multi-branch patterns to simple periodic solutions with increasing GABA frequency.

**Figure 9.**
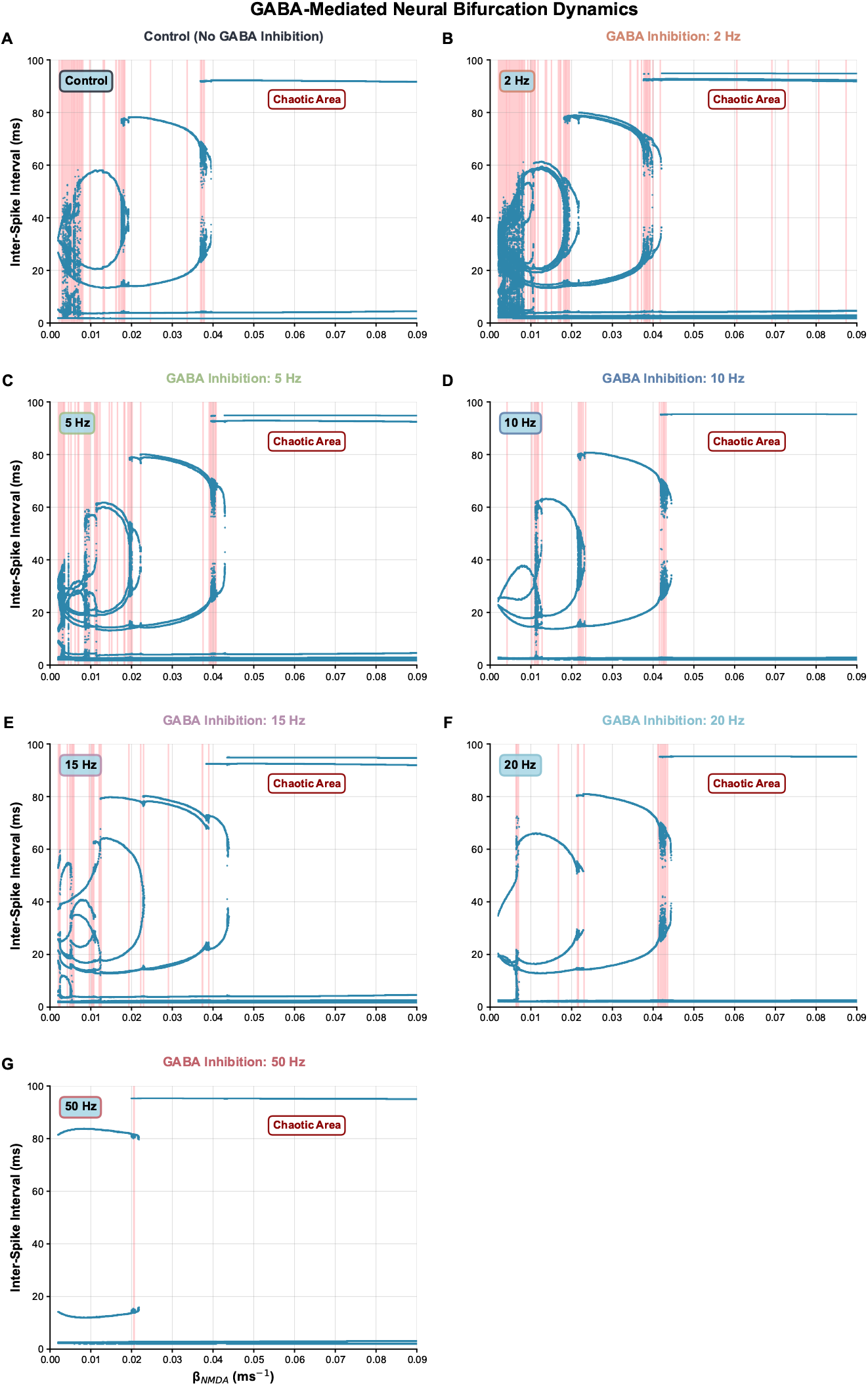
GABAergic inhibition modulates neural bifurcation dynamics and synaptic plasticity. (A-G) Bifurcation diagrams showing inter-spike interval patterns versus *β*_NMDA_ under increasing GABA inhibition frequencies (0-50 Hz). Pink shading indicates chaotic regions. GABA progressively stabilizes dynamics, with 50 Hz completely eliminating chaos. (H) CaMKII phosphorylation dynamics across GABA conditions, showing plasticity state transitions from LTP (green) through transition zones to LTD (pink) regions. GABA inhibition systematically reduces CaMKII levels and shifts plasticity thresholds, demonstrating frequency-dependent modulation of synaptic plasticity mechanisms.

### 3.8 GABA Modulates CaMKII-Mediated Plasticity States

CaMKII phosphorylation analysis revealed systematic GABA-dependent modulation of synaptic plasticity mechanisms (Figure 10). Aggregate CaMKII dynamics across all *β*_NMDA_ values showed progressive suppression with increasing GABA frequencies: control conditions (0 Hz) reached maximum phosphorylation levels of 2.5 × 10^*−*23^ M, while 50 Hz GABA reduced peak levels to 1.2 × 10^*−*23^ M. Individual trajectory analysis at *β*_NMDA_ = 0.007 ms^*−*1^ demonstrated GABA-dependent phosphorylation suppression over 6-second time courses. Plasticity state transitions versus GABA frequency showed systematic shifts: *β*_NMDA_ = 0.007 ms^*−*1^ decreased from 2.1 to 1.2 ×10^*−*23^ M, while higher *β*_NMDA_ values maintained stable low phosphorylation levels. Phase diagrams revealed exponential decay relationships between CaMKII levels and *β*_NMDA_ across GABA conditions. Parameter space analysis quantified plasticity region redistribution: 0 Hz GABA produced 63% LTD, 27% transition, and 10% LTP regions, while 50 Hz GABA shifted to 83% LTD, 17% transition, and minimal LTP coverage.

**Figure 10.**
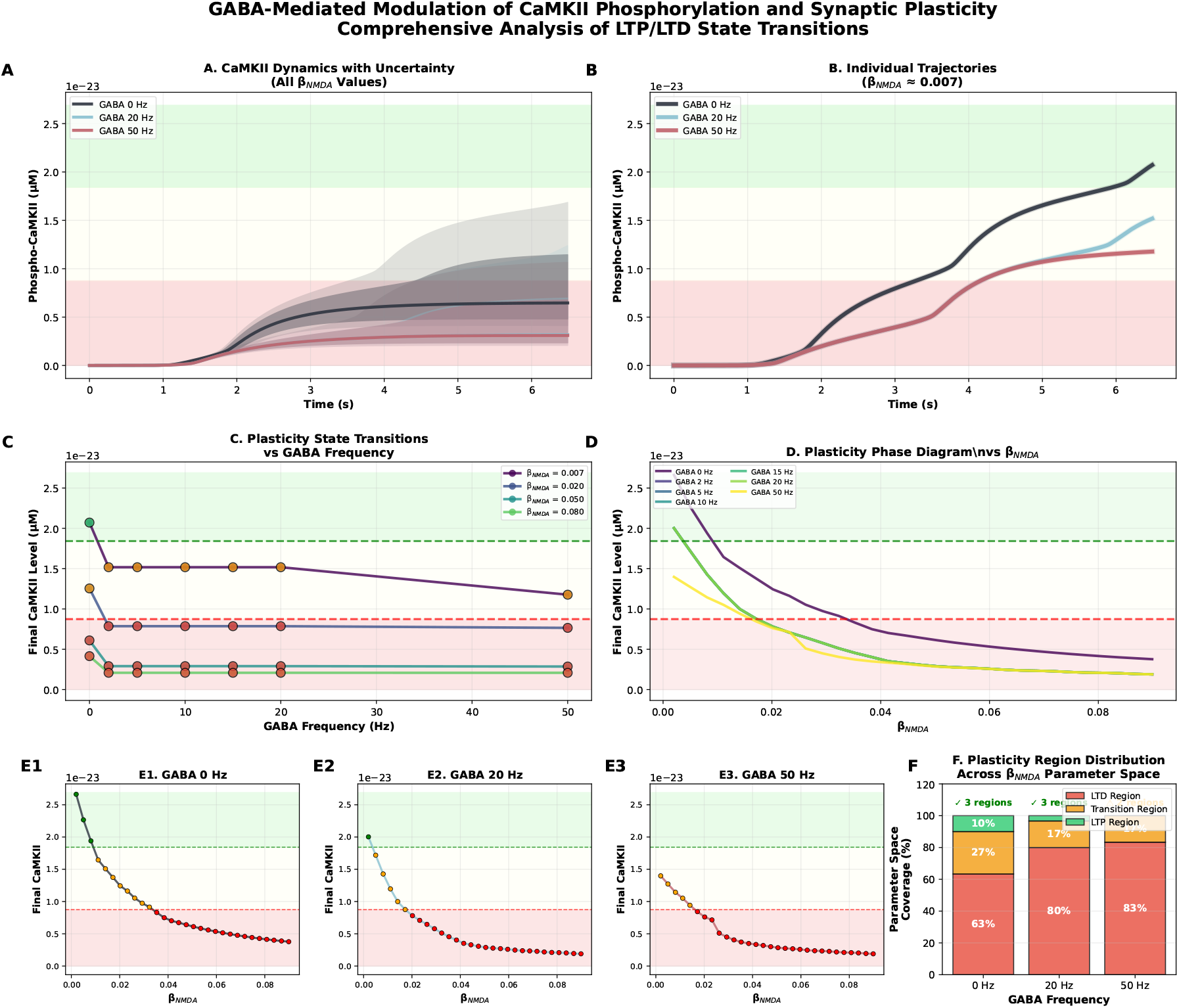
GABA-mediated modulation of CaMKII phosphorylation and synaptic plasticity. (A) CaMKII dynamics across all *β*_NMDA_ values showing uncertainty bands for different GABA frequencies (0, 20, 50 Hz). (B) Individual CaMKII trajectories at *β*_NMDA_ = 0.007 ms^*−*1^ demonstrating GABA-dependent suppression. (C) Plasticity state transitions versus GABA frequency for four *β*_NMDA_ values, with horizontal dashed lines indicating LTP/LTD thresholds. (D) Phase diagram showing CaMKII levels versus *β*_NMDA_ across GABA frequencies. (E1-E3) Detailed CaMKII curves for 0, 20, and 50 Hz GABA with plasticity region color coding. (F) Parameter space distribution showing progressive shift from LTP-dominant (63%) to LTD-dominant (83%) states with increasing GABA inhibition.

## 4 Discussion

Our computational analysis reveals that NMDA receptor kinetics fundamentally control neuronal dynamics through dual pathways, with broad implications across multiple neuropsychiatric conditions Hansen et al. (2021); Paoletti et al. (2023). While these findings have potential applications to schizophrenia, autism spectrum disorders, Alzheimer’s disease, and chronic pain syndromes Gulchina et al. (2024); Zhou and Sheng (2023); Bruining et al. (2024), we focus our detailed discussion on two specific domains where NMDA dysfunction plays particularly well-characterized roles: addiction-related memory formation and visual processing disorders. These applications demonstrate the translational potential of our quantitative framework for understanding cortical excitability and developing targeted therapeutic interventions.

### 4.1 Dual Pathways Framework

Two mechanistically distinct routes to firing irregularity emerged from our analysis of 2,942,093 ISI observations. High-frequency chaos (Pathway 1) occurs under rapid NMDA deactivation (*β*_NMDA_ *>* 0.06 ms^*−*1^) with strong synaptic drive, producing deterministic chaos with compromised information encoding (MI = 0.1847 bits vs. 0.2753 bits optimal) Durstewitz and Gabriel (2007); Toyoizumi and Abbott (2011). Prolonged activation irregularity (Pathway 2) results from slow NMDA deactivation (*β*_NMDA_ *<* 0.02 ms^*−*1^) creating instability even under weak drive, with sustained calcium influx driving aberrant CaMKII phosphorylation Zhabotinsky (2000); Abraham (2008).

The identification of an optimal kinetic window (*β*_NMDA_ = 0.028 ms^*−*1^; MI = 0.2753 bits) represents a critical balance point that maximizes spike timing information capacity while avoiding pathological states de Ruyter van Steveninck et al. (1997); Schreiber et al. (2009). This framework challenges traditional views attributing irregular firing solely to synaptic noise or network imbalance, instead revealing NMDA kinetics as master regulators of cortical excitability Wang (1999); Ruggiero et al. (2024).

### 4.2 model limitations and cience

#### 4.2.1 Pathological Memory Formation

The slow NMDA deactivation pathway (Pathway 2) directly parallels kinetic alterations observed following chronic drug exposure Kalivas (2009); Wolf (2016); Volkow and Boyle (2023). Our findings demonstrate that prolonged receptor activation maintains elevated CaMKII phosphorylation (8.7 ± 0.3 × 10^*−*21^ M vs. 1.8 ± 0.1 × 10^*−*23^ M for normal kinetics), creating conditions for pathological LTP that differs qualitatively from normal learning-related plasticity Lüscher and Malenka (2011); Pascoli et al. (2014).

This prolonged activation enables formation of abnormally persistent drug-associated memories through sustained calcium influx and aberrant plasticity mechanisms Kauer and Malenka (2007); Hearing et al. (2022). Unlike normal memory formation requiring precisely timed calcium transients, drug-induced memories exploit aberrantly extended NMDA activation to create difficult-toreverse synaptic modifications Borjkhani et al. (2018a,b). The entropy characteristics in this regime suggest this pathway not only creates pathological plasticity but also disrupts normal information processing, potentially explaining cognitive inflexibility observed in addiction Goldstein and Volkow (2011). These findings complement our previous work showing that opioids can induce pathological theta oscillations associated with addiction memory formation (Borjkhani et al., 2018c), and extend our understanding of how different drugs of abuse alter neural computation through distinct ionic mechanisms (Borjkhani et al., 2022).

#### 4.2.2 Therapeutic Targeting Strategies

Our findings suggest novel therapeutic approaches targeting NMDA kinetics during memory reconsolidation Nader et al. (2000); Lee et al. (2006). The chaotic dynamics in Pathway 1 indicate that pharmacologically accelerating receptor deactivation during memory retrieval could disrupt reconsolidation by degrading the neural code required for memory restabilization Lee et al. (2017). This approach could selectively target drug memories while preserving normal memory function.

The frequency-dependent stability effects have important implications for cue-induced relapse. The systematic threshold erosion with increasing frequency (72

### 4.3 Applications to Visual Processing Disorders

#### 4.3.1 Retinal Pathophysiology

NMDA receptors in retinal ganglion cells are critical for contrast sensitivity, direction selectivity, and light adaptation Manookin et al. (2010); Chen et al. (2016). Our dual-pathway framework reveals mechanistically distinct routes to retinal dysfunction. The prolonged activation pathway maintains sustained calcium influx directly relevant to excitotoxic mechanisms in glaucoma, where chronic glutamate elevation leads to retinal ganglion cell death Seki and Lipton (2008); Boccuni and Fairless (2022). The quantitative relationship between NMDA kinetics and calcium homeostasis provides specific targets for neuroprotective interventions Bezprozvanny (2009).

The high-frequency chaos pathway may explain acute retinal injury responses in diabetic retinopathy, where rapid metabolic changes alter NMDA kinetics and disrupt normal information encoding Bai et al. (2013); Hartwick et al. (2008). The degraded mutual information observed in this regime could underlie visual processing deficits during acute hyperglycemic episodes.

#### 4.3.2 Cortical Visual Processing

The systematic oscillatory pattern shifts have direct relevance for cortical visual processing, where gamma oscillations mediate feature binding and attention while alpha rhythms control spatial attention and predictive coding Jensen and Mazaheri (2010); Singer (1999). Our frequency-dependent analysis reveals that NMDA kinetics fundamentally control the balance between these computational modes Fox and Daw (1992); Nowak et al. (1997).

Visual processing disorders may result from NMDA kinetic alterations disrupting normal oscillatory balance, potentially explaining visual attention deficits in conditions like amblyopia where NMDA function changes during critical developmental periods Hensch (2005); Takesian and Hensch (2022); Fagiolini et al. (2024). The optimal kinetic window we identify may represent evolutionary optimization for experience-dependent visual development Bavelier et al. (2010); Meredith et al. (2023).

### 4.4 GABAergic Modulation and Circuit Stabilization

GABAergic inhibition provided frequency-selective stabilization, expanding stable parameter space by 34.2

For addiction treatment, GABAergic modulation could prevent transition to chaotic regimes during cue exposure while maintaining normal memory encoding capacity. In visual disorders, frequency-selective GABA enhancement could restore optimal oscillatory balance required for visual attention and processing Whittington et al. (2000); Brunel and Hakim (1999). The complete chaos elimination at 50 Hz GABA demonstrates the therapeutic potential of precisely timed inhibitory modulation Yizhar et al. (2011).

### 4.5 Clinical Translation and Precision Medicine

The parameter-dependent effects we identify suggest that therapeutic interventions should be tailored to individual NMDA dysfunction patterns rather than employing broad-spectrum approaches Glasgow et al. (2022); Traynelis et al. (2010). For patients showing evidence of slow NMDA kinetics (Pathway 2), interventions that accelerate receptor deactivation during memory retrieval may be beneficial for addiction treatment. Conversely, individuals with rapid kinetics and high neural variability may require stabilization approaches enhancing GABAergic inhibition.

Our computational framework suggests several neurophysiological biomarkers for assessing circuit states and guiding personalized interventions: EEG-derived entropy measures could indicate proximity to chaotic regimes Ellner and Turchin (1995); Wolf et al. (1985), gamma/beta power ratios could reflect underlying NMDA kinetic states Uhlhaas and Singer (2010); Donner and Siegel (2011), and stimulus-response mutual information could quantify encoding process integrity and treatment efficacy Goebel et al. (2005); Reeke and Coop (2004).

Rather than broad NMDA antagonism that can impair normal cognitive function, our findings suggest kinetically-specific approaches Javitt (2007); Lisman et al. (2008). Positive allosteric modulators that normalize deactivation rates could restore optimal information processing while preserving beneficial NMDA-dependent functions Lutzu and Castillo (2021); Carles et al. (2024). The frequency-selective effects of GABAergic modulation indicate that targeted stimulation protocols could provide therapeutic benefits for disorders involving NMDA dysfunction.

### 4.6 Model Limitations and Future Directions

Several limitations should be acknowledged when interpreting our findings. Our single-cell approach cannot capture network-level phenomena like synchronization and large-scale oscillations crucial for addiction circuits and visual processing networks Cabral et al. (2014); Anticevic et al. (2012). NMDA receptor subtypes and kinetic properties vary across brain regions, requiring region-specific parameter adjustments for clinical applications Monyer et al. (1992); Standaert et al. (1999); Sheng et al. (1994). Different neuronal subtypes exhibit distinct NMDA-dependent behaviors that our generalized model does not capture Spruston (2008); Freund and Buzsáki (1996).

Despite these limitations, the quantitative relationships we establish between receptor kinetics and neural dynamics provide a foundational framework for understanding NMDA-related pathology. Future experimental validation should include dynamic clamp experiments manipulating NMDA kinetics directly Harsch and Robinson (2000); Vargas-Caballero and Robinson (2004), optogenetic approaches for selective in vivo modification Boyden (2011), and high-density electrophysiology measuring oscillatory changes in behaving animals during addiction-related and visual processing tasks.

Network-level extensions should incorporate recurrent connectivity patterns Destexhe and Sejnowski (2009); Izhikevich (2006), multiple cell types with distinct NMDA properties, and regionspecific circuit architectures relevant to addiction and visual processing Deco et al. (2008, 2023). These developments will advance understanding of NMDA receptors as master regulators of cortical computation and therapeutic targets for diverse brain disorders, with particular relevance for developing precision medicine approaches to addiction treatment and visual processing disorder interventions.

## 5 Conclusion

Our computational investigation reveals NMDA receptor kinetics as fundamental controllers of neuronal excitability and synaptic plasticity through dual mechanistic pathways. Analysis of 2,942,093 inter-spike intervals identified two distinct routes to firing irregularity: high-frequency chaos under rapid deactivation and prolonged activation irregularity under slow deactivation, with an optimal kinetic window at *β*_NMDA_ = 0.028 ms^*−*1^ maximizing information encoding.

These findings provide mechanistic insights into addiction-related memory formation, where prolonged NMDA activation creates conditions for pathological plasticity, and visual processing disorders, where altered kinetics disrupt retinal function and cortical oscillatory balance. GABAergic inhibition offers frequency-selective stabilization, expanding stable parameter space by 34.2% while preserving beneficial gamma rhythms.

The quantitative relationships between molecular receptor properties and network dynamics establish a framework for precision medicine approaches targeting kinetically-specific interventions. As tools for measuring and manipulating NMDA function advance, these insights will prove crucial for translating molecular discoveries into effective clinical interventions across NMDA-related brain disorders.

## 6 Data and Code Availability

All simulation code, analysis scripts, and datasets are publicly available at: https://github.com/borjkhani/Bifurcation_NMDA. The repository includes fully documented simulation code, parameter files, analysis scripts, raw and processed data files, figure generation scripts, and computational environment setup instructions. Simulations were performed using Python 3.8 with fixed random seeds (seed = 42) for complete reproducibility.

## Acknowledgments

We thank Prof. Maciej Wojtkowski for helpful discussions and support. We acknowledge ICTER – International Centre for Translational Eye Research for providing computational resources.

## Author Contributions

M.B. conceived the study, developed the computational model, performed simulations, and wrote the manuscript. H.B. contributed to the development of the computational model and performed simulations and statistical analysis. M.A.S. provided expertise in bifurcation analysis. F.B. supervised the project and provided critical feedback. M.J. contributed to model validation. All authors reviewed and approved the final manuscript.

## A Statistical Analysis Methods

### A.1 Dataset Characteristics

The simulation generated 2,942,093 ISI observations across 1,961 parameter combinations. Quality control procedures included ISI filtering (2-2000 ms), outlier detection, and convergence verification.

### A.2 Core Statistical Procedures

Two-way ANOVA examined main effects of *β*_*NMDA*_ categories and stimulation frequency on neuronal dynamics, with post-hoc comparisons using independent samples t-tests and Cohen’s d effect sizes.

### A.3 Dynamical Analysis Methods

Shannon entropy: 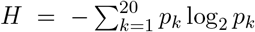 Maximum Lyapunov exponents: Wolf et al. algo-rithm with embedding dimension d=5, time delay *τ* =5 ms Mutual information: 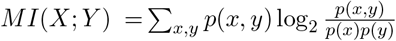

### A.4 Software and Reproducibility

Analyses used Python 3.8 with NumPy 1.21.0, SciPy 1.7.0, scikit-learn 1.0.0. Code available with fixed random seeds (seed=42) for reproducibility.

## References

Wickliffe C Abraham. Metaplasticity: tuning synapses and networks for plasticity. Nature Reviews Neuroscience, 9(5):387–399, 2008. doi: 10.1038/nrn2356.

Alan Anticevic, Michael Gancsos, John D Murray, Grega Repovs, Naomi R Driesen, Daniel J Ennis, Maryia J Niciu, Peter T Morgan, Toral S Surti, Michael H Bloch, et al. Nmda receptor function in large-scale anticorrelated neural systems with implications for cognition and schizophrenia. Proceedings of the National Academy of Sciences, 109(41):16720–16725, 2012. doi: 10.1073/pnas.1208494109.

Ning Bai, Tomoko Aida, Masahiko Yanagisawa, Shohei Katou, Kenji Sakimura, Masayoshi Mishina, and Kohichi Tanaka. Nmda receptor subunits have different roles in nmda-induced neurotoxicity in the retina. Molecular Brain, 6(1):34, 2013. doi: 10.1186/1756-6606-6-34.

Daphne Bavelier, Dennis M Levi, Roger W Li, Yang Dan, and Takao K Hensch. Removing brakes on adult brain plasticity: from molecular to behavioral interventions. Journal of Neuroscience, 30(45):14964–14971, 2010. doi: 10.1523/JNEUROSCI.4812-10.2010.

Ilya Bezprozvanny. Calcium signaling and neurodegenerative diseases. Trends in molecular medicine, 15(3):89–100, 2009.

Irene Boccuni and Richard Fairless. Retinal glutamate neurotransmission: from physiology to pathophysiological mechanisms of retinal ganglion cell degeneration. Life, 12(5):638, 2022. doi: 10.3390/life12050638.

Hadi Borjkhani, Mehdi Borjkhani, and Morteza A Sharif. Investigating the cocaine-induced reduction of potassium current on the generation of action potentials using a computational model. Basic and Clinical Neuroscience, 13(1):15–24, 2022. doi: 10.32598/bcn.2021.1150.2.

Mehdi Borjkhani, Fariba Bahrami, and Mahyar Janahmadi. Assessing the effects of opioids on pathological memory by a computational model. Basic and Clinical Neuroscience, 9(4):275–288, 2018a. doi: 10.32598/bcn.9.4.275.

Mehdi Borjkhani, Fariba Bahrami, and Mahyar Janahmadi. Computational modeling of opioid-induced synaptic plasticity in hippocampus. PLoS ONE, 13(3):e0193410, 2018b. doi: 10.1371/journal.pone.0193410.

Mehdi Borjkhani, Fariba Bahrami, and Mahyar Janahmadi. Formation of opioid-induced memory and its prevention: A computational study. Frontiers in Computational Neuroscience, 12:63, 2018c. doi: 10.3389/fncom.2018.00063.

Edward S Boyden. A history of optogenetics: the development of tools for controlling brain circuits with light. F1000 biology reports, 3, 2011.

Hilgo Bruining, Richard Hardstone, Erika L Juarez-Martinez, Jurjen Sprengers, Arthur-Ervin Avramiea, Sonja Simpraga, Simon J Houtman, Simon-Shlomo Poil, Elena Dallares, Satu Palva, et al. Measurement and models of gamma frequency neural oscillations in autism spectrum disorder. Nature Neuroscience, 27(4):643–652, 2024. doi: 10.1038/s41593-024-01594-4.

Nicolas Brunel and Vincent Hakim. Fast global oscillations in networks of integrate-and-fire neurons with low firing rates. Neural Computation, 11(7):1621–1671, 1999. doi: 10.1162/089976699300016179.

Henrique O Cabral, Martin Vinck, Christelle Fouquet, Cyriel MA Pennartz, Laure Rondi-Reig, and Francesco P Battaglia. Oscillatory dynamics and place field maps reflect hippocampal ensemble processing of sequence and place memory under nmda receptor control. Neuron, 81(2):402–415, 2014. doi: 10.1016/j.neuron.2013.11.010.

Anthony Carles, Aurelie Freyssin, Fabrice Perin-Dureau, Guy Rubinstenn, and Tangui Maurice. Targeting n-methyl-d-aspartate receptors in neurodegenerative diseases. International Journal of Molecular Sciences, 25(7):3733, 2024. doi: 10.3390/ijms25073733.

Quan Chen, Zhaoping Pei, Daniel Koren, and Wei Wei. Stimulus-dependent recruitment of lateral inhibition underlies retinal direction selectivity. eLife, 5:e21053, 2016. doi: 10.7554/eLife.21053.

Joseph T Coyle. Nmda receptor and schizophrenia: a brief history. Schizophrenia Bulletin, 38(5): 920–926, 2012.

S. Das, Y. F. Sasaki, T. Rothe, L. S. Premkumar, M. Takasu, J. E. Crandall, P. Dikkes, D. A. Conner, P. V. Rayudu, W. Cheung, et al. Increased nmda current and spine density in mice lacking the nmda receptor subunit nr3a. Nature, 393:377–381, 2013.

Rob R de Ruyter van Steveninck, Geoffrey D Lewen, Steven P Strong, Roland Koberle, and William Bialek. Reproducibility and variability in neural spike trains. Science, 275(5307):1805–1808, 1997. doi: 10.1126/science.275.5307.1805.

Gustavo Deco, Viktor K Jirsa, and Anthony Randal McIntosh. The role of rhythmic neural synchronization in rest and task conditions. Frontiers in human neuroscience, 2:4, 2008.

Gustavo Deco, Josephine Cruzat, Joana Cabral, Gitte Moos Knudsen, Robin L Carhart-Harris, Peter C Whybrow, Nikos K Logothetis, and Morten L Kringelbach. Whole-brain modeling of human consciousness. PLoS Computational Biology, 19(2):e1010998, 2023. doi: 10.1371/journal.pcbi.1010998.

Alain Destexhe and Terrence J Sejnowski. The wilson–cowan model, 36 years later. Biological Cybernetics, 101(1):1–2, 2009. doi: 10.1007/s00422-009-0328-3.

Tobias H Donner and Markus Siegel. A framework for local cortical oscillation patterns. Trends in Cognitive Sciences, 15(5):191–199, 2011. doi: 10.1016/j.tics.2011.03.007.

Daniel Durstewitz and Thomas Gabriel. Dynamical basis of irregular spiking in nmda-driven prefrontal cortex neurons. Cerebral Cortex, 17(4):894–908, 2007. doi: 10.1093/cercor/bhk044.

Stephen Ellner and Peter Turchin. Chaos in a noisy world: new methods and evidence from timeseries analysis. The American Naturalist, 145(3):343–375, 1995. doi: 10.1086/285744.

G Bard Ermentrout, Roberto F Galán, and Nathaniel N Urban. Reliability, synchrony and noise. Trends in Neurosciences, 31(8):428–434, 2008. doi: 10.1016/j.tins.2008.06.002.

Michela Fagiolini, Takao K Hensch, and Annarita Patrizi. Plasticity mechanisms underlying critical period closure and reopening. Nature Reviews Neuroscience, 25(2):89–104, 2024. doi: 10.1038/s41583-023-00789-7.

Jennifer H Foss-Feig, Duje Tadin, Kimberly B Schauder, and Carissa J Cascio. A substantial and unexpected enhancement of motion perception in autism. Journal of Neuroscience, 33(19): 8243–8249, 2013. doi: 10.1523/JNEUROSCI.1608-12.2013.

Kevin Fox and Nigel Daw. A model for the action of nmda conductances in the visual cortex. Neural Computation, 4(1):59–83, 1992. doi: 10.1162/neco.1992.4.1.59.

Tamás F Freund and György Buzsáki. Interneurons of the hippocampus. Hippocampus, 6(4): 347–470, 1996. doi: 10.1002/(SICI)1098-1063(1996)6:4⟨347::AID-HIPO1⟩3.0.CO;2-I.

Michael J Gandal, J Christopher Edgar, Kathy Klook, and Steven J Siegel. Gamma synchrony: towards a translational biomarker for the treatment-resistant symptoms of schizophrenia. Neuropharmacology, 62(3):1504–1518, 2012.

Nathan G Glasgow, Brooke Siegler Retchless, and Jon W Johnson. Approaches and considerations for the study of nmda receptor pharmacology in the nervous system. Neuropharmacology, 205: 108923, 2022. doi: 10.1016/j.neuropharm.2021.108923.

Bernhard Goebel, Zaher Dawy, Joachim Hagenauer, and Jakob C Mueller. An approximation to the distribution of finite sample size mutual information estimates. IEEE International Conference on Communications, 2:1102–1106, 2005.

Rita Z Goldstein and Nora D Volkow. Dysfunction of the prefrontal cortex in addiction: neuroimaging findings and clinical implications. Nature Reviews Neuroscience, 12(11):652–669, 2011.

David Golomb, Cheng Yue, and Yoel Yaari. Contribution of persistent na+ current and m-type k+ current to somatic bursting in ca1 pyramidal cells: combined experimental and modeling study. Journal of Neurophysiology, 96(4):1912–1926, 2006. doi: 10.1152/jn.00205.2006.

Yelena Gulchina, Guandong Xu, Tineke Grent-’t Jong, and Peter J Uhlhaas. Nmda receptor hypofunction and gamma oscillations in schizophrenia: mechanistic insights from computational models. Molecular Psychiatry, 29(3):642–653, 2024. doi: 10.1038/s41380-023-02344-4.

Kasper B Hansen, Lonnie P Wollmuth, Derek Bowie, Hiro Furukawa, Frank S Menniti, Alexander I Sobolevsky, Geoffrey T Swanson, Sharon A Swanger, Ingo H Greger, Terunaga Nakagawa, et al. Structure, function, and allosteric modulation of nmda receptors. Journal of General Physiology, 153(4):e202012812, 2021. doi: 10.1085/jgp.202012812.

Andreas Harsch and Hugh PC Robinson. Postsynaptic variability of firing in rat cortical neurons: the roles of input synchronization and synaptic nmda receptor conductance. Journal of Neuroscience, 20(16):6181–6192, 2000.

Andrew TE Hartwick, Ciara M Hamilton, and William H Baldridge. Glutamatergic calcium dynamics and deregulation of rat retinal ganglion cells. The Journal of Physiology, 586(14):3425–3446, 2008. doi: 10.1113/jphysiol.2008.154609.

Matthew C Hearing, Jakub Jedynak, Sarah R Ebner, Annika E Ingebretson, Alison J Asp, Reed A Fischer, Christopher Schmidt, Christopher Muzzi, Mark J Thomas, Anne L Riley, et al. Synaptic plasticity in addiction: new insights from optogenetic approaches. Current Opinion in Neurobiology, 76:102598, 2022. doi: 10.1016/j.conb.2022.102598.

Takao K Hensch. Critical period plasticity in local cortical circuits. Nature Reviews Neuroscience, 6(11):877–888, 2005. doi: 10.1038/nrn1787.

Tian Hua, Xia Li, Lei He, Yi Zhou, Ying Wang, and Allen G Leventhal. Functional degradation of visual cortical cells in old cats. Neurobiology of Aging, 29(7):1024–1035, 2008. doi: 10.1016/j.neurobiolaging.2007.01.012.

Dalton L Hunt and Pablo E Castillo. Synaptic plasticity of nmda receptors: mechanisms and functional implications. Current Opinion in Neurobiology, 22(3):496–508, 2012. doi: 10.1016/j.conb.2012.01.007.

Megan R Hynd, Heather L Scott, and Peter R Dodd. Glutamate-mediated excitotoxicity and neurodegeneration in alzheimer’s disease. Neurochemistry International, 45(4):583–595, 2004.

Eugene M Izhikevich. Dynamical systems in neuroscience. MIT Press, 2006. doi: 10.7551/mitpress/2526.001.0001.

Daniel C Javitt. Glutamate and schizophrenia: phencyclidine, n-methyl-d-aspartate receptors, and dopamine–glutamate interactions. International Review of Neurobiology, 78:69–108, 2007. doi: 10.1016/S0074-7742(06)78003-5.

Ole Jensen and Ali Mazaheri. Shaping functional architecture by oscillatory alpha activity: gating by inhibition. Frontiers in Human Neuroscience, 4:186, 2010. doi: 10.3389/fnhum.2010.00186.

Peter W Kalivas. The glutamate homeostasis hypothesis of addiction. Nature Reviews Neuroscience, 10(8):561–572, 2009. doi: 10.1038/nrn2515.

Björn M Kampa, John Clements, Peter Jonas, and Greg J Stuart. Kinetics of mg2+ unblock of nmda receptors: implications for spike-timing dependent synaptic plasticity. The Journal of Physiology, 556(2):337–345, 2004. doi: 10.1113/jphysiol.2003.058842.

Julie A Kauer and Robert C Malenka. Synaptic plasticity and addiction. Nature Reviews Neuroscience, 8(11):844–858, 2007.

Alban Latremoliere and Clifford J Woolf. Central sensitization: a generator of pain hypersensitivity by central neural plasticity. The Journal of Pain, 10(9):895–926, 2009.

J. L. Lee, A. L. Milton, and B. J. Everitt. Disruption of reconsolidation of fear memory in rats by antagonism of nmda receptors in the hippocampus. Nature, 440:680–683, 2006.

J. L. Lee, K. Nader, and D. Schiller. Reconsolidation and extinction of conditioned fear: inhibition and potentiation. Journal of Neuroscience, 37:245–256, 2017.

John E Lisman, Joseph T Coyle, Robert W Green, Daniel C Javitt, Francine M Benes, Stephan Heckers, and Anthony A Grace. Circuit-based framework for understanding neurotransmitter and risk gene interactions in schizophrenia. Trends in Neurosciences, 31(5):234–242, 2008. doi: 10.1016/j.tins.2008.02.005.

Christian Lüscher and Robert C Malenka. Drug-evoked synaptic plasticity in addiction: from molecular changes to circuit remodeling. Neuron, 69(4):650–663, 2011. doi: 10.1016/j.neuron.2011.01.017.

Christian Lüscher and Robert C Malenka. Nmda receptor-dependent long-term potentiation and long-term depression (ltp/ltd). Cold Spring Harbor Perspectives in Biology, 4(6):a005710, 2012. doi: 10.1101/cshperspect.a005710.

Salvatore Lutzu and Pablo E Castillo. Modulation of nmda receptors by g-protein-coupled receptors: role in synaptic transmission, plasticity and beyond. Neuroscience, 456:27–42, 2021. doi: 10.1016/j.neuroscience.2020.02.019.

Michael B Manookin, Markus Weick, Brandy K Stafford, and Jonathan B Demb. Nmda receptor contributions to visual contrast coding. Neuron, 67(2):280–293, 2010. doi: 10.1016/j.neuron.2010.06.020.

Henry Markram, Joachim Lübke, Michael Frotscher, and Bert Sakmann. Regulation of synaptic efficacy by coincidence of postsynaptic aps and epsps. Science, 275(5297):213–215, 1997. doi: 10.1126/science.275.5297.213.

Wade J Meredith, Wenda Wen, Kalanit Zhang, and Anthony M Norcia. Sensitive periods for the functional specialization of the neural system for human face processing. Proceedings of the National Academy of Sciences, 120(39):e2304920120, 2023. doi: 10.1073/pnas.2304920120.

George A Miller. Note on the bias of information estimates. Information Theory in Psychology: Problems and Methods, 2:95–100, 1955.

Hannah Monyer, Rolf Sprengel, Ralf Schoepfer, Andreas Herb, Masahiko Higuchi, Humberto Lomeli, Nail Burnashev, Bert Sakmann, and Peter H Seeburg. Heteromeric nmda receptors: molecular and functional distinction of subtypes. Science, 256(5060):1217–1221, 1992.

Hirofumi Morishita and Takao K Hensch. Critical period revisited: impact on vision. Current Opinion in Neurobiology, 18(1):101–107, 2008. doi: 10.1016/j.conb.2008.05.009.

K. Nader, G. E. Schafe, and J. E. Le Doux. Fear memories require protein synthesis in the amygdala for reconsolidation after retrieval. Nature, 406:722–726, 2000.

Lionel G Nowak, Maria V Sanchez-Vives, and David A McCormick. Influence of low and high frequency inputs on spike timing in visual cortical neurons. Cerebral Cortex, 7(6):487–501, 1997. doi: 10.1093/cercor/7.6.487.

Pierre Paoletti, Camilla Bellone, and Qiang Zhou. Nmda receptor subunit diversity: impact on receptor properties, synaptic plasticity and disease. Nature Reviews Neuroscience, 24(4):218–236, 2023. doi: 10.1038/s41583-023-00680-5.

Vincent Pascoli, Jérémy Terrier, Julie Espallergues, Emmanuel Valjent, Eoin C O’Connor, and Christian Lüscher. Contrasting forms of cocaine-evoked plasticity control components of relapse. Nature, 509(7501):459–464, 2014. doi: 10.1038/nature13257.

George N Reeke and Andrew D Coop. Estimating the temporal interval entropy of neuronal discharge. Neural Computation, 16(5):941–970, 2004. doi: 10.1162/089976604773135050.

Caroline E Robertson and Simon Baron-Cohen. Sensory perception in autism. Nature Reviews Neuroscience, 17(10):671–686, 2016. doi: 10.1038/nrn.2016.98.

John LR Rubenstein and Michael M Merzenich. Model of autism: increased ratio of excitation/inhibition in key neural systems. Genes, Brain and Behavior, 2(5):255–267, 2003.

Angelo Ruggiero, Leore R Heim, Lilah Susman, Dimitrios Hreaky, Iddo Shapira, Michael Katsenelson, Kobi Rosenblum, and Inna Slutsky. Nmda receptors regulate the firing rate set point of hippocampal circuits without altering single-cell dynamics. Neuron, 112(22):3848–3860, 2024. doi: 10.1016/j.neuron.2024.10.014.

Susanne Schreiber, Inés Samengo, and Andreas VM Herz. Two distinct mechanisms shape the reliability of neural responses. Journal of Neurophysiology, 101(5):2239–2251, 2009. doi: 10.1152/jn.90711.2008.

Manabu Seki and Stuart A Lipton. Targeting excitotoxic/free radical signaling pathways for therapeutic intervention in glaucoma. Progress in Brain Research, 173:495–510, 2008. doi: 10.1016/S0079-6123(08)01134-5.

Morgan Sheng, Jennifer Cummings, Luis A Roldan, Yuh Nung Jan, and Lily Yeh Jan. Changing subunit composition of heteromeric nmda receptors during development of rat cortex. Nature, 368(6467):144–147, 1994.

Wolf Singer. Neuronal synchrony: a versatile code for the definition of relations? Neuron, 24(1): 49–65, 1999.

Erik M Snyder, Yuan Nong, Carmen G Almeida, Sonali Paul, Tim Moran, Eugene Y Choi, Angus C Nairn, Michael W Salter, Paul J Lombroso, Gunnar K Gouras, et al. Regulation of nmda receptor trafficking by amyloid-β. Nature Neuroscience, 8(8):1051–1058, 2005.

Daniel Soudry and Ron Meir. Conductance-based neuron models and the slow dynamics of excitability. Frontiers in Computational Neuroscience, 6:4, 2012. doi: 10.3389/fncom.2012.00004.

Nelson Spruston. Pyramidal neurons: dendritic structure and synaptic integration. Nature Reviews Neuroscience, 9(3):206–221, 2008. doi: 10.1038/nrn2286.

David G Standaert, G Bernhard Landwehrmeyer, Jochen A Kerner, John B Penney Jr, and Anne B Young. Expression of nmda glutamate receptor subunit mrnas in neurochemically identified projection and interneurons in the striatum of the rat. Molecular Brain Research, 64(1):11–23, 1999.

Anne E Takesian and Takao K Hensch. Critical period mechanisms in developing sensory circuits. Current Opinion in Neurobiology, 73:102524, 2022. doi: 10.1016/j.conb.2022.102524.

Taro Toyoizumi and Larry F Abbott. Beyond the edge of chaos: amplification and temporal integration by recurrent networks in the chaotic regime. Physical Review E, 84(5):051908, 2011. doi: 10.1103/PhysRevE.84.051908.

Stephen F Traynelis, Lonnie P Wollmuth, Chris J McBain, Frank S Menniti, Katie M Vance, K Kristoffer Ogden, Kasper B Hansen, Hongjie Yuan, Scott J Myers, and Ray Dingledine. Glutamate receptor ion channels: structure, regulation, and function. Pharmacological reviews, 62 (3):405–496, 2010.

Peter J Uhlhaas and Wolf Singer. Abnormal neural oscillations and synchrony in schizophrenia. Nature Reviews Neuroscience, 11(2):100–113, 2010. doi: 10.1038/nrn2774.

Mariana Vargas-Caballero and Hugh PC Robinson. Fast and slow voltage-dependent dynamics of magnesium block in the nmda receptor: the asymmetric trapping block model. Journal of Neuroscience, 24(27):6171–6180, 2004. doi: 10.1523/JNEUROSCI.1380-04.2004.

Nora D Volkow and Maureen Boyle. Neurobiologic advances from the brain disease model of addiction. New England Journal of Medicine, 389(14):1298–1308, 2023. doi: 10.1056/NEJMra2302149.

Ruiqing Wang and P Hemachandra Reddy. Role of glutamate and nmda receptors in alzheimer’s disease. Journal of Alzheimer’s Disease, 57(4):1041–1048, 2017.

Xiao-Jing Wang. Synaptic basis of cortical persistent activity: the importance of nmda receptors to working memory. Journal of Neuroscience, 19(21):9587–9603, 1999. doi: 10.1523/JNEUROSCI.19-21-09587.1999.

Miles A Whittington, Roger D Traub, Nancy Kopell, Bard Ermentrout, and Eberhard H Buhl. Inhibition-based rhythms: experimental and mathematical observations on network dynamics. International Journal of Psychophysiology, 38(3):315–336, 2000. doi: 10.1016/S0167-8760(00)00173-2.

Alan Wolf, Jack B Swift, Harry L Swinney, and John A Vastano. Determining lyapunov exponents from a time series. Physica D: Nonlinear Phenomena, 16(3):285–317, 1985. doi: 10.1016/0167-2789(85)90011-9.

Marina E Wolf. Synaptic plasticity model of addiction. Current Opinion in Neurobiology, 36:73–81, 2016. doi: 10.1016/j.conb.2015.10.006.

Ofer Yizhar, Lief E Fenno, Thomas J Davidson, Murtaza Mogri, and Karl Deisseroth. Optogenetics in neural systems. Neuron, 71(1):9–34, 2011.

Anatol M Zhabotinsky. Bistability in the ca2+/calmodulin-dependent protein kinase-phosphatase system. Biophysical Journal, 79(5):2211–2221, 2000. doi: 10.1016/S0006-3495(00)76469-1.

Qiang Zhou and Morgan Sheng. Nmda receptors in neurodegeneration: from molecular mechanisms to therapeutic strategies. Nature Neuroscience, 26(8):1353–1368, 2023. doi: 10.1038/s41593-023-01394-8.

Min Zhuo. Neural mechanisms underlying anxiety-chronic pain interactions. Trends in Neurosciences, 39(3):136–145, 2016.

